# Mucosal ecosystems of the ileum and colon shape the therapeutic response in Crohn’s Disease

**DOI:** 10.1101/2025.03.26.645552

**Authors:** Ran Zhou, Bingqing Xie, Christopher Weber, Joel Pekow, Candace M. Cham, Jianqiao Liu, Yu Zhao, Jason Koval, Eugene B. Chang, Anindita Basu

## Abstract

Many Crohn’s disease patients fail to respond to biologic therapies targeting canonical immune pathways, and the mechanisms underlying treatment non-response remain poorly understood. Integrating single-cell and spatial transcriptomics, we generated a comprehensive mucosal atlas of 380,000 cells spanning the terminal ileum and ascending colon of healthy donors and Crohn’s disease patients receiving biologics. We revealed site-specific programs centered on TNF-expressing macrophages embedded within epithelial-stromal networks: follicle-associated enterocytes dictated immune activation in the ileum, while FAP-expressing stromal cells served this role in the colon. Co-expression network analysis further identified lineage-specific programs associating non-response, including a novel epithelial marker BPIFB1 independent of known genetic risk loci. Co-expression marker Z-scores significantly differed between responders and non-responders in published datasets. Together, these findings define a non-responder-specific epithelial-stromal-immune ecosystem that sustains chronic inflammation, providing a framework for stratifying treatment resistance and informing therapeutic strategies in Crohn’s disease and other barrier organ disorders.

## INTRODUCTION

Crohn’s disease (CD) is an immune-mediated disorder affecting the digestive tract, especially the small and large intestines (*1*). Approximately 10 million patients have been diagnosed with CD worldwide, and more than half of this population require life-long medical intervention (*1*). Dysregulated immune responses are central to CD, involving both myeloid (neutrophil, macrophage, dendritic cell) and lymphoid (CD4^+^ helper T, regulatory T, cytotoxic CD8^+^ T, plasma B cell) populations (*2*). These cells accumulate in the local environment, engaging activities mediated by proinflammatory molecules including tumor necrosis factor (TNF) α, interleukin (IL)-12/23, integrin α4(β7) and Janus kinases (JAK 1/3) (*1, 2*). Accordingly, monoclonal antibodies targeting these molecules constitute first-line biologic treatments for CD.

Despite the clinical success of biologics, over one third of patients fail to respond to these treatments (non-responders), and a subset of initial responders lose efficacy over time (1, *2*). Persistent inflammation in these non-responders— both primary and secondary— poses a major clinical challenge with significant implications for public health outcomes and healthcare costs (*1*). Furthermore, this response heterogeneity suggests that disease-maintaining mechanisms extend beyond canonical immune pathways due to the complexity of the intestinal microenvironment (*3–7*).

A prevailing model of CD pathogenesis proposes that increased epithelial barrier permeability permits the influx of microbial antigens from the lumen—an environment uniquely burdened by a dense and diverse microbiota (*2, 4*). This influx triggers potent immune activation within the underlying lamina propria (*2, 4, 6*). Although epithelial barrier dysfunction is a hallmark of CD, the mechanisms allowing and maintaining a “leaky” epithelium remain poorly defined. It is unclear whether epithelial dysfunction reflects passive damage or an active, regulated cell state shift. Notably, both the intestinal epithelial barrier and mucosal immune systems are highly adaptive to environmental pressures, raising the possibility that chronic disease drives maladaptive epithelial-mediated programs that sustain the disease (2, *4, 6*).

CD non-responders harbor a constrained inflammatory environment in which immune actions are continuously suppressed, yet inflammatory flares persist (1, 6). This setting provides a unique opportunity to interrogate disease mechanisms that are unmasked by therapeutic intervention. We therefore hypothesize that, under disease and treatment pressures, epithelial cells adopt functions that engage conventional immune cells, amplify immune activation, and destabilize intestinal integrity.

To test this hypothesis, we focused on intestinal mucosa from non-responsive CD patients, including specimens with ileitis and colitis, to capture site-specific disease features.

Using single cell transcriptomics, we constructed comprehensive cellular networks for human mucosa from the terminal ileum (TI) and ascending colon (AC), encompassing both CD and healthy donors. We further resolved cell-cell neighbors and signals using spatial transcriptomics. This integrative analysis reveals an epithelium-stroma–macrophage network that is selectively activated in non-responsive CD. The epithelium and its stromal neighbors exhibit immune-engaging programs and spatial proximity to inflammatory macrophages, showing the hypothesis that the epithelium contributes to chronic inflammation has merit. Together, these findings highlight epithelial–immune-stromal interactions as a potential driver of treatment-refractory CD and provide a resource for further mechanistic and therapeutic exploration.

## RESULTS

We profiled and analyzed 85 intestinal mucosal biopsies from 17 non-IBD and 29 CD donors (Fig. 1A, Table S1). CD samples were classified by patient clinical response and by inflammatory status: inflamed (Inf), adjacent-to-inflamed (Adj, histologically uninvolved) and uninvolved (Uni) (Methods). Non-responders (n=17, Inf and Adj) included recurrent or refractory patients primarily receiving anti-TNFα biologics and anti-IL12/23 therapies. CD donors in clinical remission (Uni) were categorized as responders. Post-treatment, patients – regardless of responses - exhibited change in disease involvement (Fig. S1A). Although our cross-sectional design precludes temporal inference, these changes reflect therapy-associated mucosal remodeling. Our cohort is among the first few to be deliberately powered for treatment non-responders, enabling direct mucosal comparison across healthy controls, responders and non-responders under different biologic pressures.

**Figure 1.**
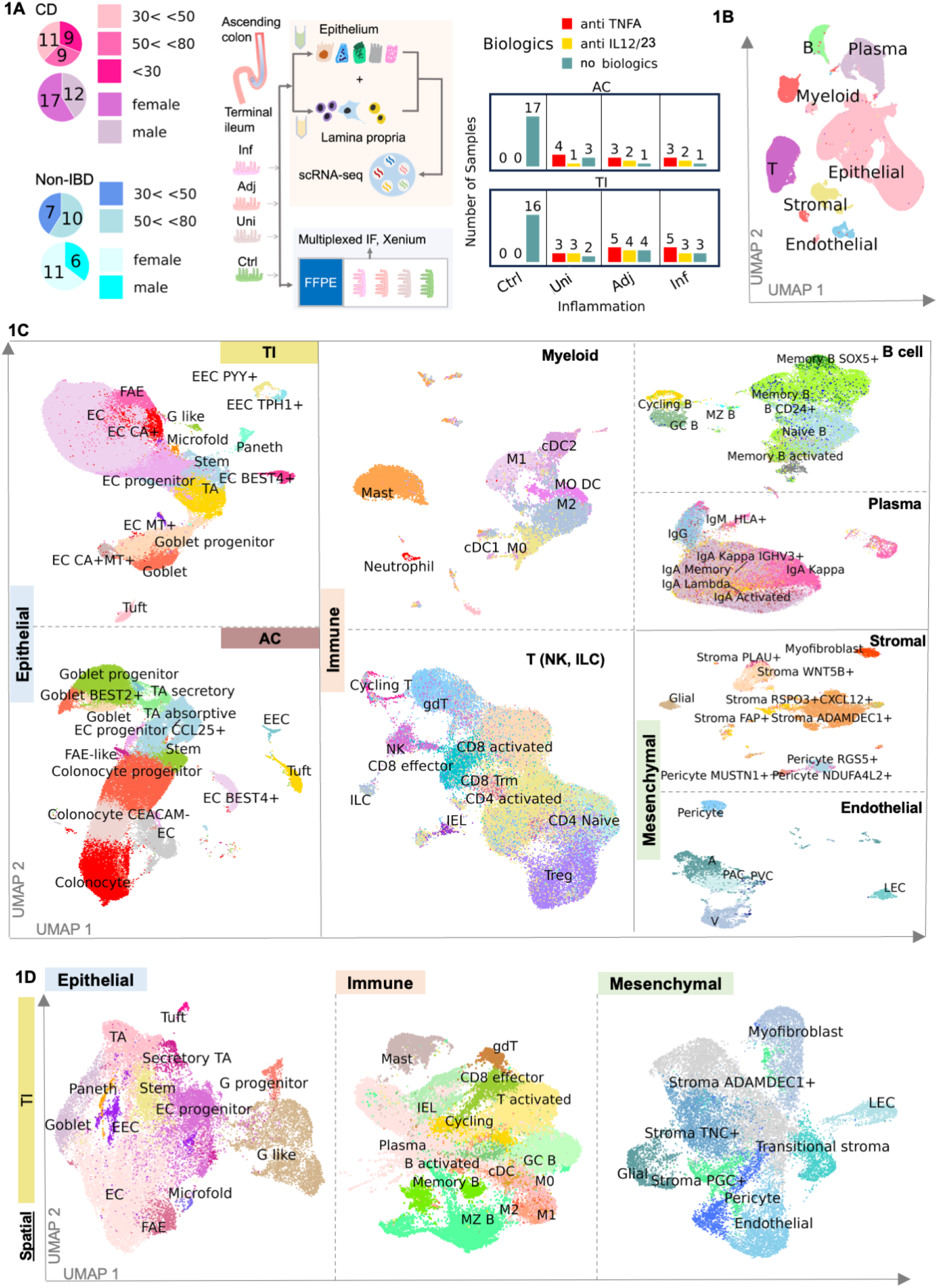
Single cell atlas of TI and AC biopsies from non-IBD control donors and CD patients. A. Study design integrating scRNA-seq, Spatial transcriptomic and IF. Left, donor ages: middle, workflow schematic; right, donor treatments. B. Compartmental-level census across all samples. C. Cell type census on all cells in UMAPs. Lineage: epithelial (TI and AC), immune (T/NK/ILC, B, Plasma and Myeloid), and mesenchymal (stroma, endothelium). D. TI-specific lineage and cell type census from spatial data (Xenium).

### 1. Mucosal atlas reveals known and previously unrecognized cellular states

In total, we recovered 383,522 high-quality cells with three defined lineages: epithelial, immune and mesenchymal from sc transcriptomes. We categorized lineages into compartments: epithelial from the terminal ileum (TI) or ascending colon (AC), lymphoid and myeloid, stromal and endothelial (Fig. 1B).

We identified 79 cell types (Methods: Table 1) exhibiting transcriptomic consistency with previous studies, including rare types, e.g., Paneth, tuft and microfold cells, (*4,6–8*) (Fig. 1C, S1B). In this study, we highlight a few cell types that contribute to the ecosystem specific to non-responders: 1) macrophages, including proinflammatory M1-like macrophage (referred to as M1, expressing *TNF*), and M2-like alternative macrophage (referred to as M2) (*9*); 2) follicle associated enterocyte (FAE), an ileum specific, specialized enterocyte expressing *CCL20* and *IL15*. FAE cells were identified by transcriptome sharing features with both enterocytes and microfold cells, yet distinctly lacked GP2 expression — a canonical marker for mature microfold cells (*4,7,8*); 3) gastric chief-like cells—G-like cells—an epithelial metaplasia (Fig. 1C, S1B), and 4) site-specific stromal cells including stroma PLAU+ in the ileum and stroma FAP+ in the colon (Fig. 1C, S1B). We did not recover any enteric neurons after the enzymatic dissociation.

**Table 1.**
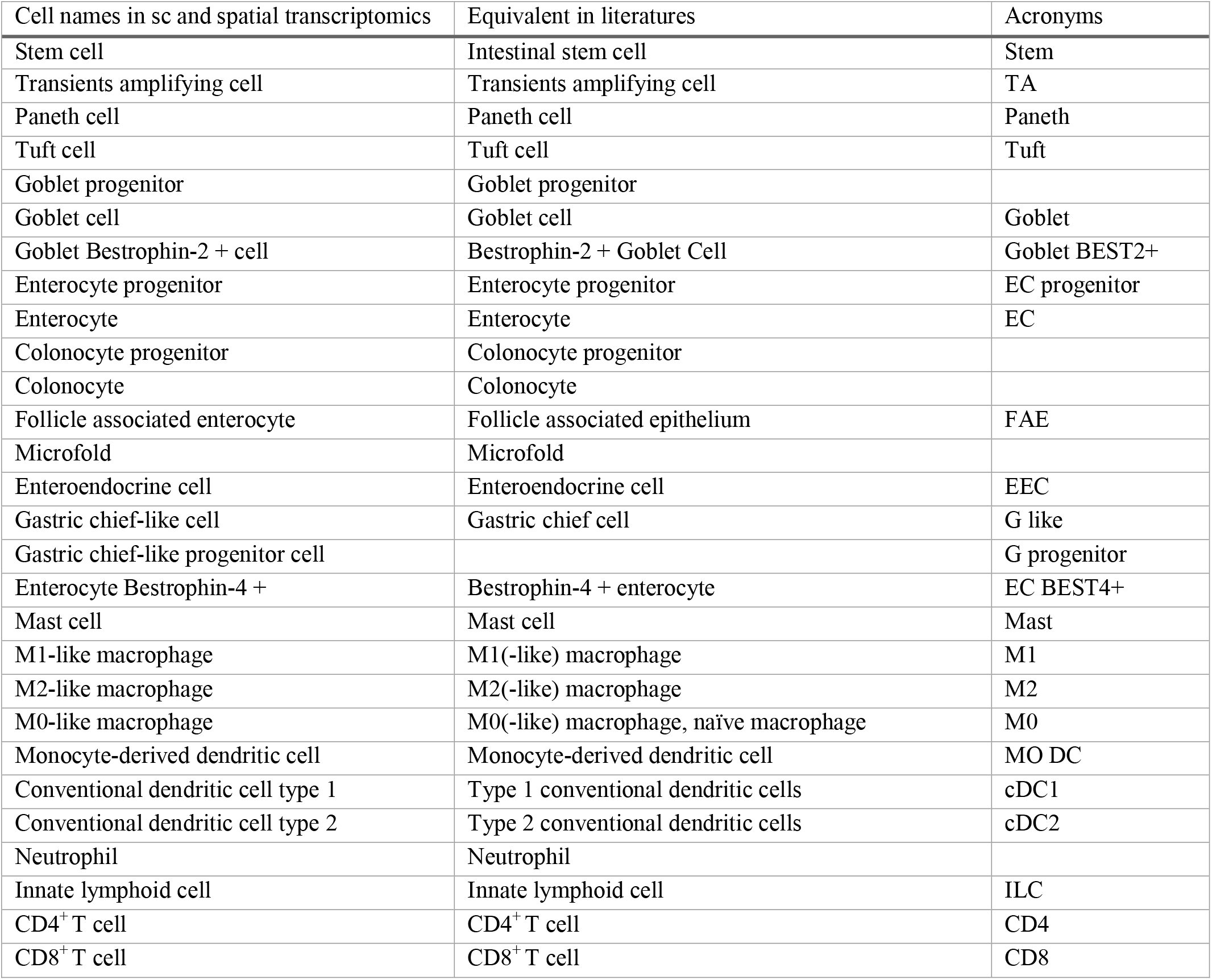
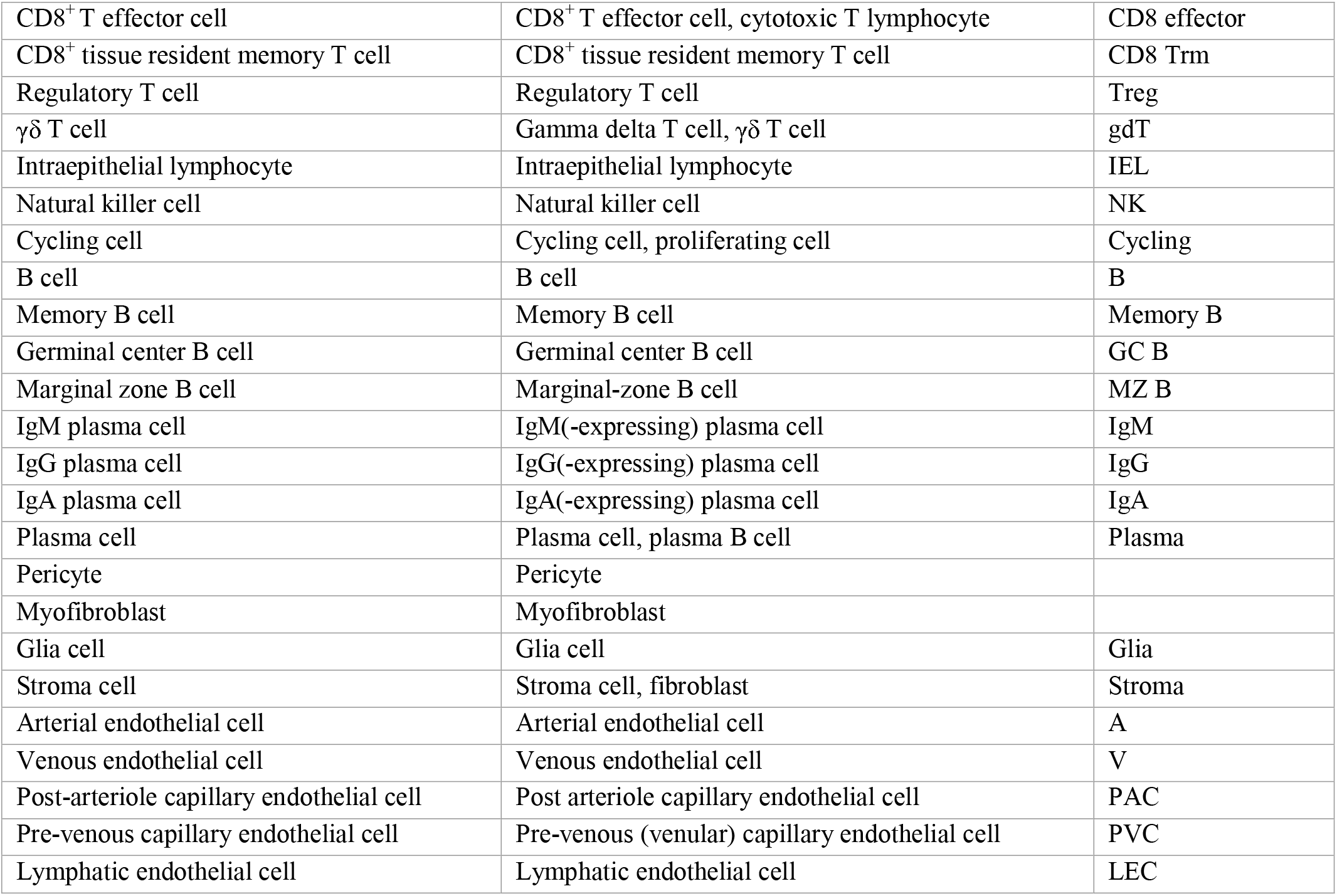
Cell names in sc and spatial transcriptomics.

To continue the cell profiling in the intricate microenvironment, we applied spatial transcriptomics (Xenium) on ileal mucosa covering phenotypes—Ctrl, Adj, Inf, Inflamed with epithelial metaplasia (Inf_metaplasia). We recovered 86,313 high-quality cells, of which many cell types similar to those in sc transcriptomes, in the spatial atlas (Fig. 1D). The spatial atlas had different reportage on cell identity, reflecting variation in sampling and transcript coverage. For example, cycling B and T cells were annotated collectively as cycling; meanwhile, the spatial atlas enabled further delineation of several subtypes including G-like progenitor, stroma PGC+, stroma TNC+, and secretory TA (Fig. 1D). Although all the annotations were influenced by assay complexity (scRNA-seq ∼18000 genes vs spatial ∼450 genes) and clustering resolution, the concordance between the two atlases supported downstream analysis for cellular crosstalk and neighborhood in these tissues.

Prior to analyses, we assessed the experimental and biological covariates in donors that could contribute to transcriptional variability across donors (Fig. S1C). Mucosal classifiers (Ctrl, Uni, Adj, Inf) and administered biologics (treatment) were the dominant factors driving intra-compartmental variance (Fig. S1C). The mucosal transcriptome is predominantly shaped by disease state and treatment response, validating the analytical framework focus on the cell state change and interaction shift.

### 2. A non-responder–specific epithelial–immune–stromal network underlies persistent inflammation

We first examined the cell types and programs that are associated with persistent inflammation in non-responsive CD.

#### 2.1. Non-responder mucosa exhibits site-specific inflammatory cellular organization

Although mucosa from different patient groups had similar lineage compositions (Fig S2A), M1 expansion is the most prominent feature in non-responders, providing a cellular basis for the sustained inflammatory environment that underlies treatment failure. (Fig. 2A). Notably, differences associated with treatment response in ileum are featured by FAE. The disparity in cellular composition across Ctrl, Uni (responder) and Inf (non-responder) were accompanied by the transcriptomic changes (Fig. 2B). In both regions, non-responders highly express *IL1B* and *IGHG3* in myeloid and plasma, as well as *LCN2* in epithelium. Although pan markers for epithelium (Fig. 2B; *4, 10*) remain consistent across inflammation status, non-responders have site-specific epithelial expressions—*CCL25, GATA4, REG1A* in TI, *CXCL13* in AC (Fig. 2B; *11-13*). The lineage-specific gene upregulation with increasing inflammation severity suggests that the mucosal inflammatory response is not driven by a single cell population but is coordinated across lineages —indicating an ecosystem-level activity rather than cell type-level phenomenon.

**Figure 2.**
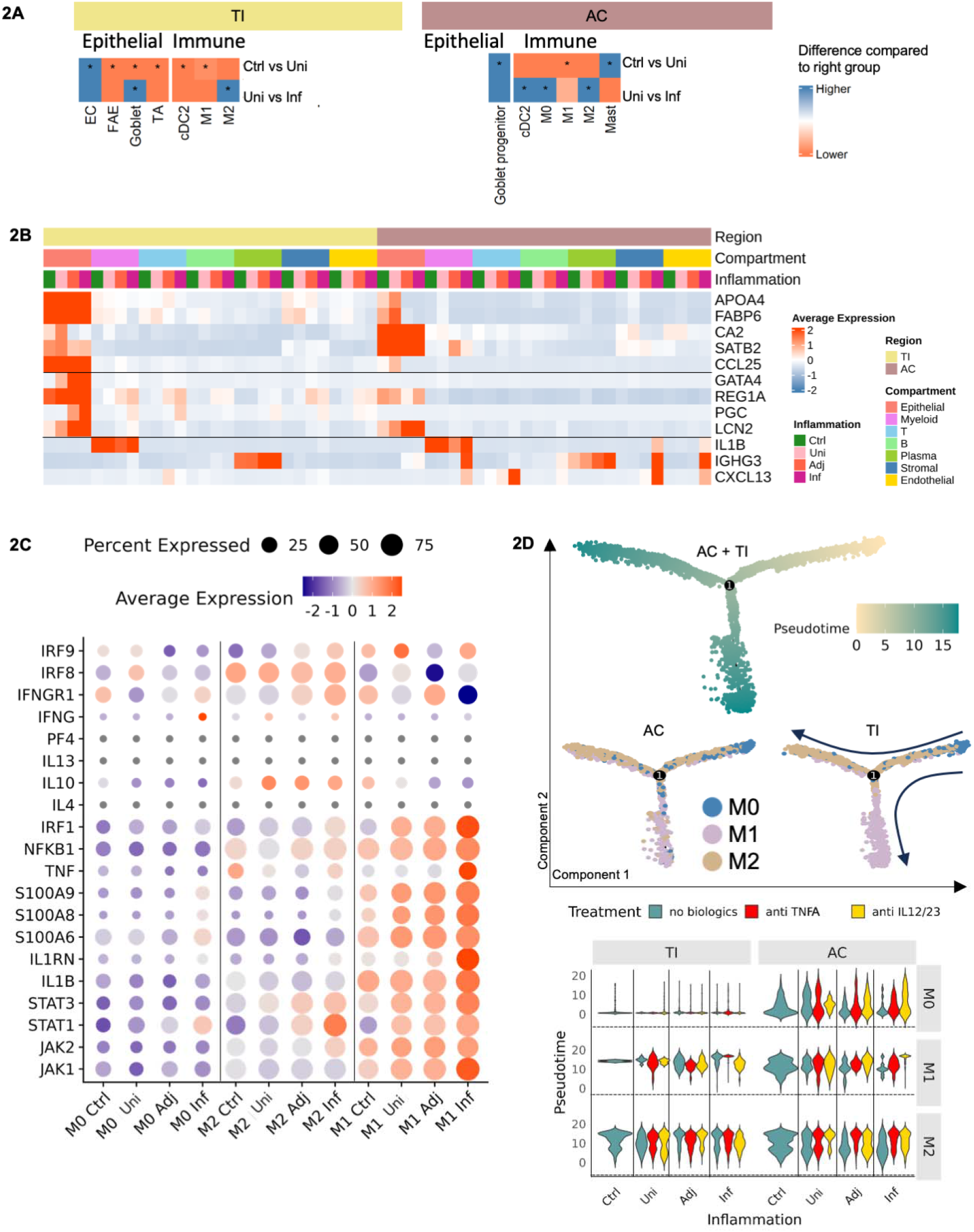
Inflammation-associated changes in cell composition and gene expression A. Significant changes of cell composition in TI and AC. Difference of means shown, red indicates increase in second group, blue in the first group; Wilcoxon test; *, P<0.05. B. DEG at compartment level across inflammation classification. C. DEG in macrophages. X-axis, macrophage from different inflammation classification; y-axis, genes. D. Macrophage trajectory. Top, DDRtree trajectory of macrophages; bottom, pseudo-time distribution under different treatments along the imputed trajectory.

Below, we outline site-specific patterns in cellular transcription and interactions across all the lineages that illustrate inflammatory networks in non-responders.

##### 2.1.1 TNF-expressing inflammatory macrophages are a hallmark of inflamed non-responder mucosa

Macrophages have diverse roles: pro-inflammatory M1 enhance immune reactivity, while anti-inflammatory M2 enhance immune tolerance and tissue repair (*14*). Overall macrophage presence increases in the mucosa with inflammation (Fig. S2B, Table S1). Each macrophage subtype exhibits distinct expression patterns. M1 showed elevated expression of proinflammatory genes (*IL1B, IRF1, S100A6, S100A8, S100A9*), and contribute to inflammation directly by upregulating *TNF* and *IL1RN* (Fig. 2D). Polarization genes (*JAK1, JAK2, STAT1* and *STAT3*) (*15–17*) were widely expressed in M1, but only expressed in a small fraction of M0 and M2 (Fig. 2C, Table S2). M2 showed minimal change in abundance, as well as expression of anti-inflammatory cytokines (*IL4, IL10, IL13*; *14*). However, interferon signaling (*IFNGR1*, *IRF1, IRF8, IRF9*) were upregulated in the M2, suggesting activation of proinflammatory pathways (Fig. 2C, Table S2; *17*). To examine the relations within these subtypes, we performed pseudotime trajectory analysis to infer transcriptional progression and state transitions of macrophages based on transcriptomic similarity. Combining with DEGs, pseudotime trajectory analysis indicates that the coexistence of M0, M1, and M2 within the mucosa reflects a spectrum of activation states with distinct functions: M0 as a plastic precursor, M2 as an anti-inflammatory population with a subset that serves as emerging precursors, and M1 as the dominant pro-inflammatory effector — whose TNFα expression are the molecular targets of current biologic therapies. Pseudotime variations in these subtypes under different treatments were intestinal location-dependent, except for M2. M0 and M1 dynamics highlight their sensitivity to local microenvironmental cues. The stability of M2 pseudotime profiles across intestinal locations, treatment groups, and patient groups suggests M2 polarization is a conserved response largely independent of treatment. The distinct pseudotime positioning of M1 suggests it represents a terminal inflammatory state, that polarization is established and may be hard to transition to another state. The divergence of M1 pseudotime between responders and non-responders, and across treatment types, raises the possibility that the trajectory of macrophage polarization — not just its endpoint — may distinguish patients’ responses to current biologics.

We next assessed pathway activity by comparing *TNF* and *IL12/IL23* pathway module scores. M1, MO DC and ILC consistently shows the highest *TNF* scores, irrespective of inflammation, sampling site or treatment, while *IL12* and *IL23* scores remained high in NK (Fig. S2B). B lymphocytes and plasmablasts showed no expression of all these pathways. These results indicate that immune cells are the common effector of cytokine-driven inflammation in both small and large intestines.

Among the common effectors in non-responders, macrophages emerge as critical with population expansion. Their polarization dynamics suggest they do not operate in isolation. This prompted us to investigate the upstream cellular partners that orchestrate and sustain macrophage-driven inflammatory activity in the mucosal ecosystem.

##### 2.1.2 Myeloid crosstalk defines inflammatory cellular neighborhoods

To further examine cells that shape the ecosystem, we employed ligand-receptor (LR) analysis to infer cell-cell interactions. These interactions revealed activities involving all three lineages, with increasing connectivity and complexity in the order: Ctrl< Responder < Non-responder (Table S3). From our previous analysis, macrophages are central to pro-inflammatory activities. Thus, in the imputed interactions, we focused on the cells that signaled to macrophage and dendritic cells. These interactions are presented as networks, with nodes representing cell types and edges indicating LR pairing and their diversities. In ileum, M2 primarily interacted with epithelial cells and stroma ADAMDEC1+ in Ctrl and these interactions were similar in responders. In non-responders, macrophage interactions extended to FAE, cDC2, IgA plasma, and endothelial cells in ileum, while Stroma ADAMDEC1+ selectively interacted with M1 in non-responder (Fig. 3A). Interactions in AC were more complicated, involving two stromal cells, stroma ADAMDEC1+ and stroma WNT5B+ (Fig. 3B). In both Ctrl and non-responder groups, M2 and MO DC received signals from these stromal cells. Furthermore, M1 extended their interaction to stroma FAP+, a cell type enriched in non-responders. Evidently, stromal cells consistently participated in the network, forming the interaction core in AC.

**Figure 3.**
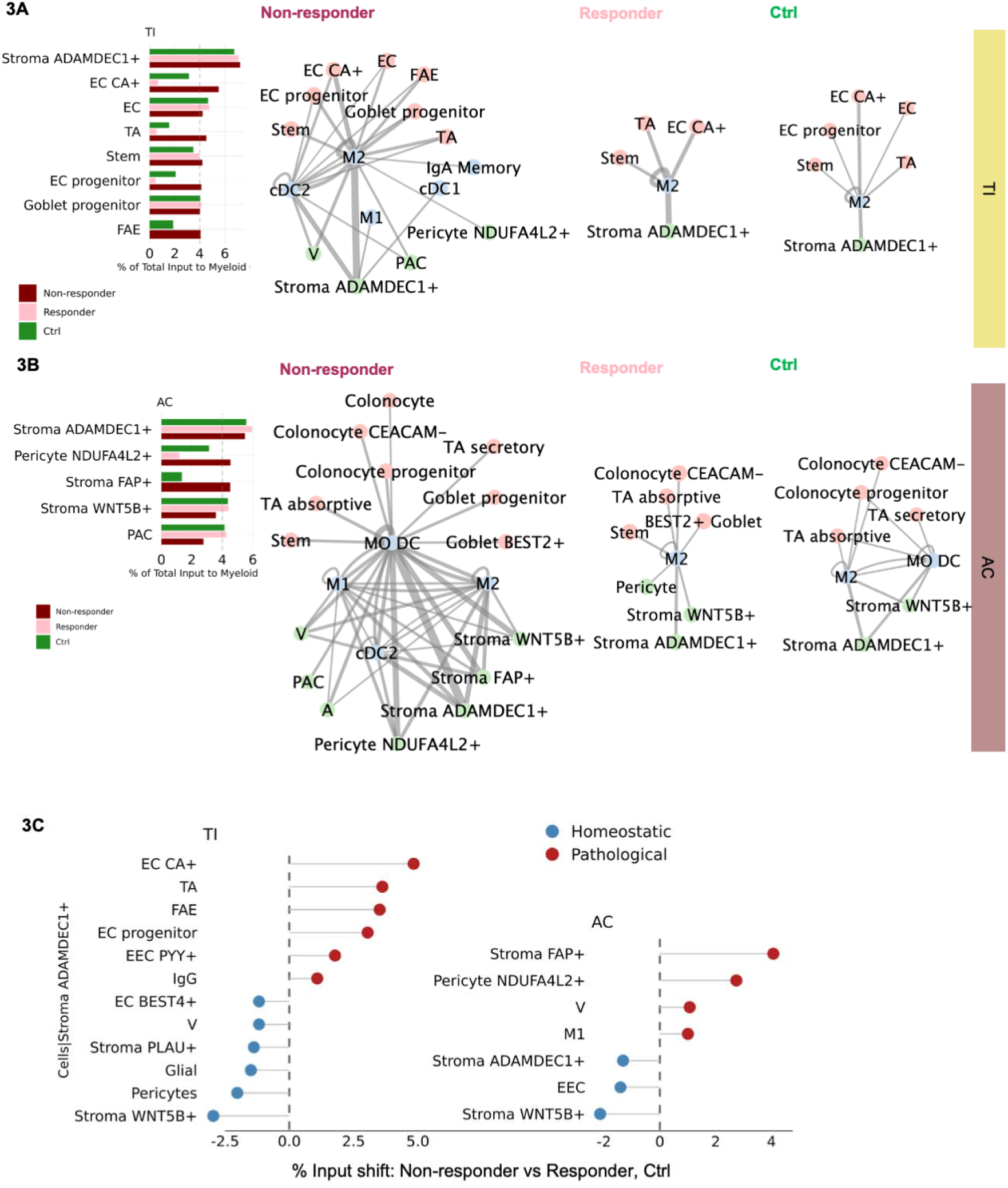
Inflammation associated changes in cell-cell interactions. A. LR analysis in TI. Left, cell types that contribute to LR connections in the interaction network (myeloid cells = R-expressing). Right, interaction map (myeloid cells = R-expressing) in Ctrl, responder and non-responder. B. LR analysis in AC. Left, cell types that contribute to LR connections in the interaction network (myeloid cells = R-expressing). Right, interaction map (myeloid cells = R-expressing) in Ctrl, responder and non-responder. Node color: lineage (blue – epithelial, pink – immune, green– mesenchymal); edge thickness – normalized LR pairs. C. Leading cell types that contribute to LR connections in the interaction network (Stroma ADAMDEC1+ = R-expressing).

To complement the LR-lead crosstalk, we analyzed the cellular neighbors in a subset of ileum mucosa. First, we compartmentalized the mucosal sections by 17 gross domains across non-IBD donors (Ctrl) and non-responders (Fig. S2C). Within each domain, we identified the most prevalent cell-cell spatial neighbors. Notably, the most common cell neighbors with macrophages remain consistent across Ctrl and non-responder groups, including epithelial and B/plasma cells, except for metaplasia-specific cells (G progenitor) (Fig. S2C). This indicated that epithelial cells were irreplaceable components assembling the inflammatory ecosystem, while M1 served as critical effectors in inflamed mucosa.

#### 2.2 Stromal cells organize inflammatory cell interactions

Stromal cells are composed by fibroblasts (as ‘Stroma marker+’) and shape the extra-cellular environment in the intestines (Fig. 1C, S1B; *18,19*). To identify the dominant upstream signaling sources, we quantified each cell type’s relative contribution to the total LR-connecting network. This identified the stromal ADAMDEC1+ as the primary “signal broadcaster” to the myeloid networks (Fig. 3A, 3B). This stromal subtype was highly conserved across ileum, colon and patient groups, showing transcriptomic stability (cosine similarity >0.9). Such high similarity suggests a stable, site-independent stromal identity rather than a localized, context-specific response.

Applying the same logic to identify the drivers upstream of Stroma ADAMDEC1+, we found interactions specifically increased in non-responders, and they showed divergence between ileum and colon. In the ileum, the signals were dominated by TA, EC progenitor, EC CA+, FAE and EEC PYY+, among which FAE significantly increased in non-responders; conversely, the signals in the colon were driven by Stroma FAP+, Pericyte NDUFA4L2+, V, and M1, with stroma FAP+ and M1 increased in non-responders (Fig. 2A, 3C). This site-specific variation resonates with the distinct function and structure of the gut, including fibroblasts typically associated with colonic IBD (*20,21*). We further inspected molecular components in the interactions involving stroma ADAMDEC1+ (Fig. S2D). The distinct LR interactions that differentiate between patient groups, were TGFB1/TGFBR1 in Ctrl, and TNF/TNFRSF1A in non-responder. The cell neighbors specific to non-responders were TA, EC progenitor, FAE and G-like cells. Overall, these results indicate stroma ADAMDEC1+ transmit signals from epithelium to myeloid.

In all, the non-responder mucosal ecosystem was characterized by a denser intercellular network — expanded in both breadth and depth of signaling. Within this network, the conserved stroma ADAMDEC1+ emerged as the dominant upstream broadcaster to macrophages across both sites, itself receiving input from site-specific driver cells — FAE in the ileum and stromal subtypes in the colon. This hierarchical architecture identifies the stromal-macrophage axis as a conserved upstream vulnerability, with site-specific entry points, that motivate deeper characterization of these site-specific upstream populations.

#### 2.3 Specialized epithelial states mediate ileal inflammation in non-responders

The epithelium serves as a barrier between the environment and the host and is often disrupted in diseases (*22*), showing significant changes in cell composition and functions across disease conditions. We established that these cells, particularly FAE, contribute to non-responder inflammatory ecosystem. Thus, we further interrogated these special epithelial cells.

FAE and progenitors (stem, TA) significantly increased in the order Ctrl to Inf, while EC (the largest epithelial subset) decreased (Fig. 4A). Despite cytokines and chemokine expression (Fig. S1B), FAE also expressed known proinflammatory markers *LCN2, NOS2, DUOX2* (LND) (*23*), with a subset of other subtypes (Fig. S2E). Non-responders upregulated programs including regeneration (*REG1A, CD74*) (*24,25*), barrier function (*GATA4, JAK2*) (*26*), interferon signaling (*IFITM3, DMBT1*) (*27*) and oxidative stress (*DUOX2, NOS2*) (*28*) (Fig. 4A). Genes related to immune modulation, were primarily expressed by FAE (Fig. 4A, Table S4). Transcription factors (TF) associated with TNF signal, also shifted in concert with inflammation, from FOS in Ctrl to IRF1 in Inf (Fig. 4A, S3A, Table S4). These functions were complemented by upregulation of genes in GWAS risk loci (*IRF1, HSP90B1, HLA-DB1, HIF1A*) (*29*). In contrast, Ctrl functions are mainly unrelated to inflammation, while FAE still contributes to immune modulation (Fig. S3A, Table S4).

**Figure 4.**
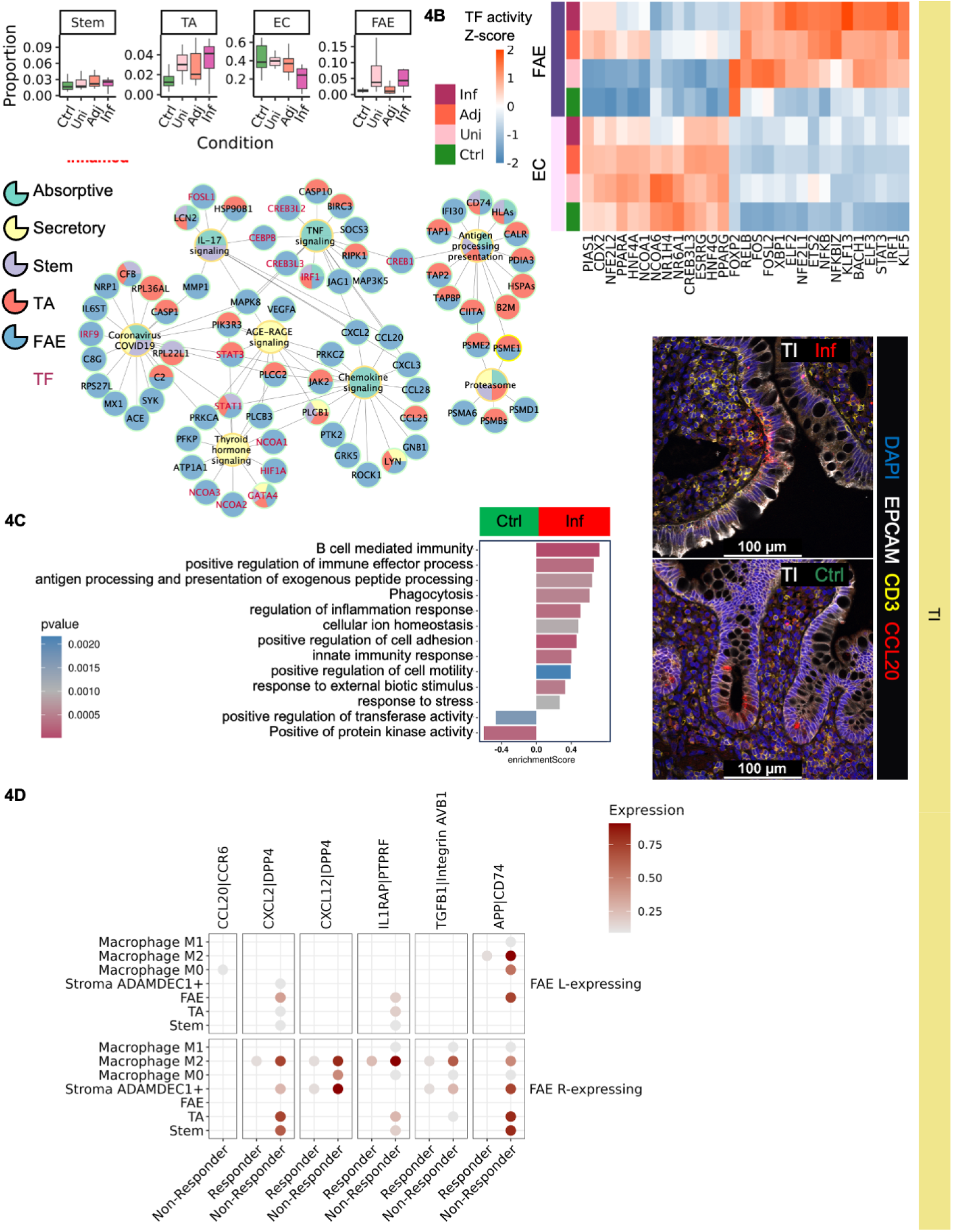
Inflammation associated epithelial programs. A. TI epithelial cell composition and function. Top, cell composition across inflammation classification. Bottom, GSEA in epithelial cells from Inf TI. Central node, enriched functions enriched; peripheral nodes, highly DE genes within the function. Node and wedge color indicate function and cell type attribution; wedge area reflects cell-type contribution. B. Predicted TF activity distinguishes EC and FAE across inflammation classification. C. Functions in Inf FAE. Left, GSEA comparison Ctrl and non-responder. Right, IF. FAE marker, CCL20 protein. D. Key FAE interaction in responder and non-responders.

##### 2.3.1 Follicle associated enterocytes activate inflammatory programs in the ileum

We used downstream gene expression-based TF activity estimation to infer cell-type characteristics of FAE, comparing them to the major absorptive cell—EC (Fig. 4B). Proinflammatory (*IRF1, NFKB, NFKBIZ*) (*30*) and survival (*KLF13*) (*31*) factors were specific to FAE and exhibit the highest activity in non-responders. FAE also displayed activities associated with epithelial-mesenchymal transition (EMT)—*BACH1, ETS2, XBP1, KLF5* (*32–34*), whereas EMT is repressed in EC via *CDX2* expression (*35*). EC also expresses *HNF4A/HNF4G,* maintaining homeostasis (*36*). The functions of non-responder FAE, in agreement with TF activity, include antigen processing, phagocytosis and signaling to immune cells (Fig. 4C, Table S4). Early FAE progenitors (stem, TA) were also enriched for immune modulation, including positive regulation of leukocyte cell-cell adhesion (GO:1903039), CCR chemokine receptor binding (GO:0048020) and negative regulation of leukocyte mediated cytotoxicity (GO:0001911) (Fig. S3B, Table S4). These progenitors exhibited increased abundance in non-responders, like FAE (Fig. 4A, S4B).

Our data also showed extensive interactions between FAE and other cells that adapt to inflammation and treatment. For instance, the location of CCL20 protein shifted from the apical side of FAE (4/4 tissues) without T cell colocalization, to the basal side in FAE (3/4 sections), where CD3 T cells colocalize (Fig. 4C), further suggesting FAE-immune axes in non-responders. Across patient groups, FAE received epithelial renewal signals (*WNTs, BMPs*), expressed by stem, TA (Table S4; *37*). Non-responder FAE also showed interactions with non-epithelial cells: with macrophage, and stroma ADAMDEC1+ via IL1 signal (*CXCL2/12, IL1RAP*), along with *CXCL2* signaling to early progenitors (*38*) (Fig. 4D, Table S3). Furthermore, macrophage, stroma ADAMDEC1+ and TA expressed anti-inflammatory *TGFB1* in non-responder, likely counteracting FAE damage (Figure 4D; *39*). Last, FAE expressed various signals to enhance immune activity, including *CCL20* and *APP,* which may recruit macrophage and lymphoid cell to the mucosa (Fig. 4D; *40, 41*).

Together, these findings establish FAE as a functionally programmed epithelial population in non-responders — expanding in abundance, acquiring immune-regulatory transcriptional identity, and spatially repositioning to direct inflammatory recruitment toward the lamina propria. Through LR communication with stroma ADAMDEC1+ and macrophages, FAE serve as the epithelial trigger of the upstream inflammatory hierarchy. Their expansion and spatial reorientation in non-responders, identifie FAE as a previously underappreciated yet pivotal orchestrator of treatment-resistant mucosal inflammation in CD.

##### 2.3.2 Epithelial metaplasia is associated with inflammation

We identified G-like cells in pyloric gland metaplasia in one non-responder patient (*42*), as evidenced by multiple assays (Fig. S3C, S3D). G-like cells in ileum were expressing *CCL2, CLL3, PGC* transcripts and OLFM4 and PGC (progastricsin) proteins, within the glands in the lamina propria (Fig. S3C). The G-like cells were wired to proinflammatory functions via *IRF1*, *JAK2* and *DUOX2* (Fig. S3D; *43*), akin to FAE exhibiting immune modulation functions (Fig. S3D).

##### 2.3.3 Patchy inflammatory lesions in non-responders involve coordinated multiple-cellular circuits

In CD, patients often have heterogeneous inflammation, with non-inflamed tissue interspersed between inflamed areas, resulting in patchy lesions (*44*). Although epithelial damage does not always coincide with transmural damage (*45*), we sought to characterize this patchiness by mapping coherent changes in cell composition and transcription for key cell types as well as function circuits in non-responders. In TI, we identified FAE-macrophage-stroma ADAMDEC1+. FAE show incremental increases in abundance, cytokine/chemokine expression (*CCL20, CCL25*, *IL15*) and TF expression (*IRF1, NFKB, NFKBIZ*), in the order Ctrl < Responder (Uni) < Non-responder (Adj, Inf) (Fig. 4B, Table S4). As the inflammatory effector, M1 abundance is positively correlated with neutrophils (Pearson correlation r =0.46, p<0.05) and inflammation classification (Spearman correlation rho = 0.435, p<0.05). Stroma ADAMDEC1+ actively engages with FAE, macrophage, cDCs in the inflamed mucosa. Collectively, these TI cells produce persistent proinflammatory signals despite treatment. In AC, neutrophils are absent in sc transcriptome atlas (Table S1), while M1 and stromal subpopulations drive inflammation: stroma FAP+ are positively correlated with inflammation classification (Spearman correlation rho = 0.475, p<0.05). While stromal FAP+ was identified as the upstream driver in the colon, their further characterization was limited by the small number of colonic non-responder in the cohort (n=3), precluding robust downstream analysis.

### 3. Co-expression networks capture transcriptional programs associated with chronic inflammation and treatment response

To complement the cellular interaction network from sc and spatial transcriptomes, we applied co-expression network analysis to capture sample-level gene coordination across the tissue — providing a lineage-level molecular perspective independent of cell-type definitions.

#### 3.1 Co-expression networks differ between the terminal ileum and ascending colon, but are conserved within response groups

Distinct from DEG, Co-expression-based inflammatory genes (*CoInfSig*) represented gene-gene association across different cells, linking gene co-occurrence and inflammation classifications (Fig. S4A, S4B) (*46*). We reconstructed a co-expression network for each inflammation classification to identify *CoInfSig* patterns. In both sites, we observed the network complexity and heterogeneity increased monotonically: Ctrl<Responder (Uni)<Non-responder (Adj, Inf) (Table S5). To enhance the generalizability of *CoInfSig*, we conducted down-sampling permutations (Fig. 5A, 5B). Down-sampling permutation testing identifies a minimal set of co-expression markers robust across varying sample sizes — prioritizing analytical confidence over signature breadth. This approach served as a proof of concept for rigorous marker selection in cohorts where sample size is a constraint. The genes with optimal generalization in the TI are *BPIFB1*, *MUC6, PGC* (n ≤5) (Fig. 5A, epithelial), and *S100A8, S100A9, IL1RN, FCGR2A* (n ≤ 5), (Fig. 5A, immune). In contrast, *FOLH1, WARS* (n>12) for epithelial, and *WDFY3, C15orf48, AP002582.*1 (n>12) for immune, did not demonstrate generalization (Fig. 5A). In AC, extended *CoInfSig* in immune demonstrated generalization (*S100A8, S100A9, IL1RN, FLT1, LIMK2, DENND5A, PPIF, ACSL1, ITGAX*, n≤5). These *CoInfSig* genes, regardless of generalizability, are associated with intestinal functions, e.g., digestion (*PGC, GIP, GATA4, KCNIP4*) (*47*), and inflammation modulation (*S100A8, S100A9, CXCL9*) (*48*).

**Figure 5.**
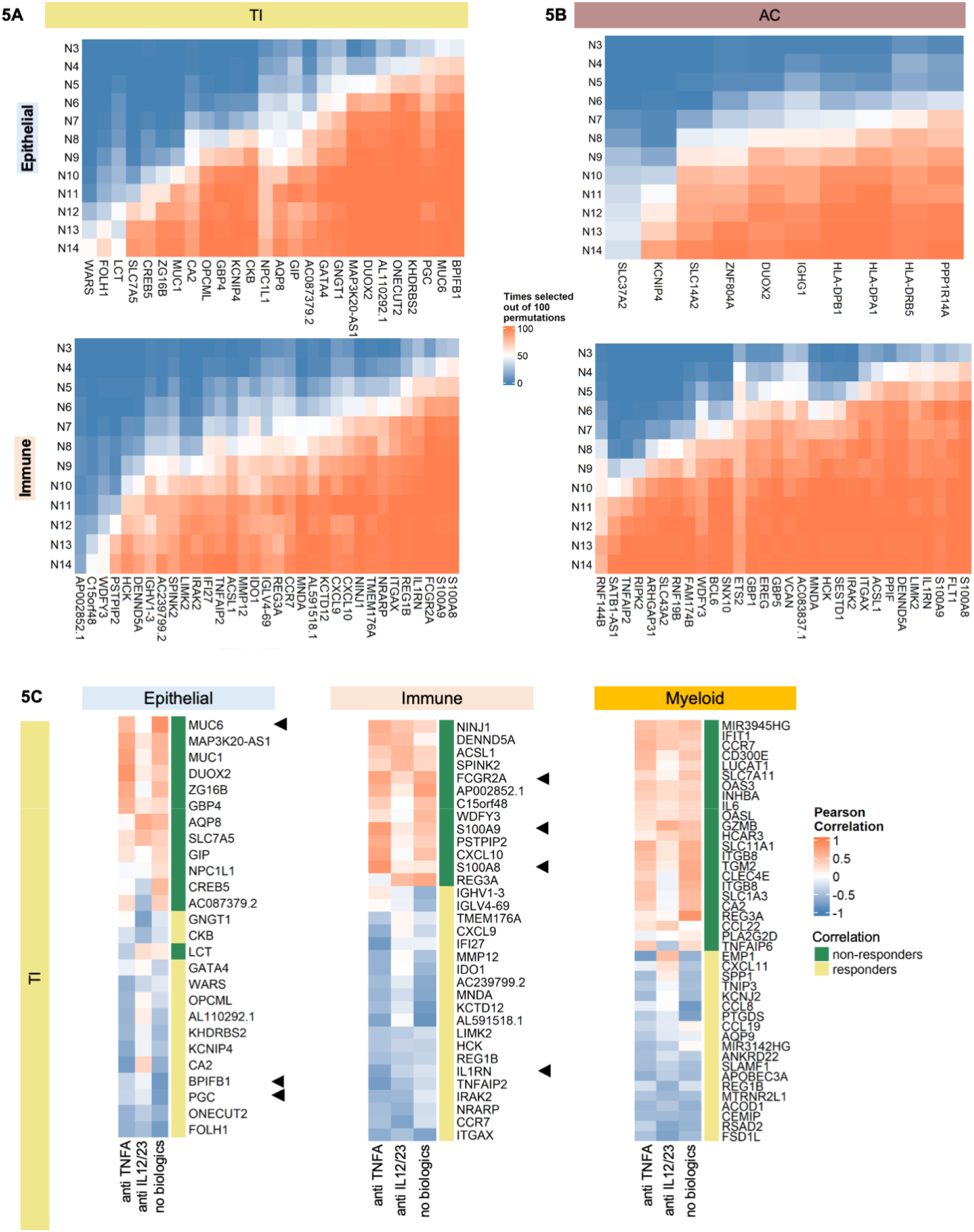
Co-expression network analysis in TI and AC. A. Reproducibility of *CoInfSig* in varied sample sizes in TI. Top, epithelial. Bottom, immune. B. Reproducibility of *CoInfSig* in varied sample sizes in AC. Top, epithelial. Bottom, immune. C Pearson correlation between the inflammation and gene loadings across biologic treatment in **TI**, Side bar indicates correlation direction. Left, epithelial; middle, immune; right, myeloid. Arrows indicate *CoInfSIg* with optimal generalization (*n* ≤ 5)

#### 3.2 Co-expression-derived gene programs stratify samples by biologic treatment and inflammatory state

To extract the gene features that undernote tissue response to biologics, we correlated a subset of tensor components (TC) with responses for the treatment (Fig. 5C). The leading genes in TCs were lineage specific. The lineage signatures for non-responders to different biologics were similar: *epithelial - MUC1, DUOX2, GBP4*, immune – *FCGR2A, S100A8, S100A9,* myeloid- *IL6, INHBA*—M1 polarization genes (*49*). A few exceptions, that are biologic-specific: epithelium -*CA2,* immune - CXCL9, myeloid - EMP1, were signatures in anti-IL-12/IL-23 non-responder. In contrast, *GIP, AC087379.2, TNGAIP6* were signatures shared by anti-TNFα and no biologics groups (treated by other immune modulators) in non-responders (Fig. 5C).

With all the signature genes identified by DE analysis, co-expression analysis and previous report on UC - *UCSig* (*6*), we graded the samples by the gene expressions (Fig. S4C) to assess the potential utilization as disease biomarkers. Notably, TI DEG expressions distinguish the samples across inflammation classification the most. In contrast, AC DEG expressions showed minimal difference across inflammation. *UCSig* expressions displayed differences in immune lineage in both sites. *CoInfSig* showed an incremental change in epithelial and immune cells from both sites. Additionally, we assessed these gene expressions in different treatment groups: the expression patterns across inflammation are not significantly affected by biologics (Fig. S4D).

#### 3.3 Co-expression-derived markers differentiated external samples by donor response to anti-TNF***α*** therapy

To explore the usability of co-expression markers outside our cohort, we scored samples by *CoInfSig* expression, across published datasets including longitudinal samples from the same donors with clinical assessment for anti-TNFα responses (*50, 51*). The *CoInfSig* markers that survived our generalization tests are listed as follows: TI epithelial (*BPIFB1, MUC6, PGC*), TI immune (*S100A8, S100A9, IL1RN, FCGR2A*), AC epithelial (*PPP1R14A, HLA-DRB5*) and AC immune (*S100A8, S100A9, IL1RN, FLT1, LIMK2, DENND5A, PPIF, ACSL1, ITGAX*).

The external cohorts were split into two major groups based on the sampling sites (TI and AC). Within each site, samples were further grouped by their timing of sampling (pre and post to administered biologic therapy) and clinical diagnoses (healthy and CD in TI, healthy, CD and UC in AC).

The power of differentiating responders and non-responders using the same set of *CoInfSig* differs between bulk transcriptomes and pseudo bulk lineage transcriptomes aggregated from single cells. When applying TI *CoInfSig* on TI bulk transcriptomes, the scores differentiate CD responder and non-responder before biologic therapy using epithelial markers (Fig. S5A) and distinguish pre and post therapy in CD and UC responders using immune markers (Fig. S5B). Yet, similar trends are not completely conserved in pseudo bulk lineage transcriptomes. Similar patterns are observed in AC, where epithelial AC *CoInfSig* scores differ between CD responders and non-responders before biologic (Fig. S5A), and immune AC *CoInfSig* scores distinguish pre and post in CD and UC responders (Fig. S5B). Because of the shared genes in TI and AC immune *CoInfSig,* the differentiating power for inflammation activity is conserved across sites and IBD subtypes (CD and UC; Fig. S5B). Overall, differentiating biologic response by epithelial *CoInfSig* was site-specific (Fig. S5A) and similar differentiation by immune *CoInfSig* was inflammation-related (Fig. S5B), with both better perform in bulk transcriptomes.

The distinct *CoInfSig* markers between epithelial and immune lineages parallel the hierarchical signaling architecture in the cellular interaction network — where epithelial cells serve as upstream driver and immune cells as downstream effector of mucosal inflammation. This positional distinction suggests that epithelial and immune markers may capture different aspects of treatment response: epithelial markers potentially reflecting upstream activity as resistance, while immune markers indexing the severity of downstream inflammatory activity.

## DISCUSSION

Chronic inflammation in CD drives cumulative tissue damage, leading to intestinal strictures, obstruction, fistula formation, and repeated surgical interventions (*1*). Although biologic therapies targeting canonical immune pathways improve disease management, a substantial fraction of CD patients experience relapse or refractory to treatments. This therapeutic non-response exposes a critical gap in our understanding of how inflammatory programs adapt under immune-targeted pressure (*1, 2, 5, 8*). Here, we show that non-responders in CD exhibit a site-specific, self-reinforcing ecosystem, in which epithelial, immune, and stromal compartments coordinate to sustain mucosal inflammation.

By integrating single-cell and spatial transcriptomics across small and large intestines, we demonstrate that TNF-expressing macrophage acts as central inflammatory hubs in non-responders (*6, 52*). However, their accumulation and activation are not autonomous. Rather, macrophages are embedded within structured peri-epithelial microenvironments scaffolded by non-immune cell types. This organization distinguishes non-responders from responders and healthy controls, suggesting that therapeutic resistance arises from tissue-level rewiring rather than isolated immune dysregulation. Inflammatory macrophages thus appear necessary but insufficient to program the local tissue state driving pathogenesis.

In the ileum, FAEs emerges as a key epithelial partner in sustaining macrophage-driven inflammation. FAEs normally interface with immune cells to support epithelial barrier, yet in non-responders, they adopt proinflammatory programs that trigger macrophage activation. This activity is associated with FAE accumulation and reprogramming, supporting a model in which epithelial cells actively sustain inflammation rather than serving as passive targets of damage (*53*). Elevated DUOX2 expression and reactive oxygen species in FAE suggest a mechanism by which epithelial stress reinforces macrophage activation and promotes epithelial-mesenchymal transition (*54, 55*). Together, these findings position the ileal epithelium as a dynamic regulator of inflammatory activity, supporting a broader principle in which epithelial plasticity contributes to the maintenance of inflammation under therapeutic pressure.

In contrast, colonic inflammation in non-responders is organized through a distinct stromal architecture. Here, stromal FAP+ cells, rather than epithelial subtypes, form the primary inflammatory interface with macrophages. The involvement of stroma ADAMDEC1+ — previously implicated in mucosal inflammation and malignancy (*6, 56, 57*)—supports a broader role for stroma shaping immune niches under disease and therapeutic stress. These findings emphasize that persistent inflammation in CD is locally encoded, with different cellular components assuming dominant regulatory roles. Such specialization may represent a general feature of barrier tissue, in which distinct non-immune compartments dominate inflammatory control depending on local structure and function. This principle has implications for tissue-level regulation in other chronic inflammatory conditions and barrier organs.

Beyond cell-type-centered networks, co-expression analysis reveals coordinated transcriptional programs within lineages. These programs are conserved within patient groups yet diverge sharply between responders and non-responders — reinforcing non-response as a distinct disease state. Using this framework, we identify epithelial markers associated with inflammation, including *BPIFB1,* which encodes a bacterial binding protein, not implicated in IBD GWAS loci (*29,58*). The emergence of BPIFB1 as a robust epithelial marker suggests that transcriptional states shaped by chronic inflammation and treatment pressure extend beyond known genetic risk architecture, implicating epithelial-microbial interactions as a contributing dimension of treatment resistance. While biologic-specific resistance markers such as *CA2* and *GIP* emerged from our analysis, limited sample sizes preclude definitive conclusions and highlight the need for larger, longitudinal cohorts encompassing different biologic therapies. Furthermore, epithelial markers distinguished responders and non-responders in CD at the pre-treatment phase in a site-specific manner, while immune markers tracked inflammation change over the course of therapy in responders in both UC and CD (*51*). This pattern, that epithelial and immune markers capture distinct aspects of treatment response, supports the usability of co-expression markers for clinical stratification, though the trend was less consistent in single cell data (*50*). Importantly, this approach may offer a generalizable framework for defining tissue-level inflammatory states and their drivers beyond CD.

Several limitations should be acknowledged. The cross-sectional design restricts causal inference regarding whether epithelial and stromal reprogramming precedes or follows immune activation. Longitudinal sampling before and after therapeutic intervention will be required to resolve the temporal hierarchy of these maladaptation. Nonetheless, the consistency of site-specific inflammatory architectures across non-responders supports the existence of stable, maladaptive tissue states that are not readily reversed by current biologic therapies.

In summary, our study reframes biologic non-response in CD as the persistence of a reconfigured mucosal ecosystem rather than incomplete immune suppression. By demonstrating how epithelial, stromal, and immune cells cooperate in a site-specific hierarchy to sustain mucosal inflammation, we provide a mechanistic framework for treatment resistance that extends beyond single-target biologic paradigms. Epithelial and stromal states — sitting upstream of the inflammatory macrophage programming — emerge as candidates for patient stratification and therapeutic targets, complementing existing immune-directed approaches. More broadly, this work illustrates how tissue ecosystem organization shapes chronic inflammation, with implications for understanding treatment resistance across other immune-mediated barrier tissue diseases.

## MATERIALS AND METHODS

### Ethical Statement

Mucosal biopsies were collected under the protocol 15573A approved by Institutional Review Board at the University of Chicago. Written consent was acquired prior to endoscopy. The endoscopic procedures were conducted by licensed gastroenterologists in compliance with relevant regulation. All donor information was anonymized and de-identified for data and sample collections.

### Sample collection by colonoscopy

The healthy donors were recruited using the following criteria: no history of IBD, no ongoing colitis or intestinal infections, no intestinal malignancies, no history of autoimmune disease or under immune modulated medications, and free of family history in autoimmune disorders, or intestinal malignancies. CD patients were grouped by physicians using colonoscopy and biologic administration history: responder– patients who show no symptoms or endoscopic, histological inflammation after receiving biologics; non-responder– patients with endoscopic inflammation and histological inflammation, regardless of symptom, after receiving biologics. Metadata for donors is provided in Table S1. Mucosal biopsies were classified by the physician and the pathologist into four categories: non-disease control (Ctrl); CD uninvolved, free of active inflammation, from treatment responders (Uni); CD inflamed, with active inflammation, from treatment non-responders (Inf); and uninvolved sites adjacent to inflamed CD lesions (Adj). All the tissues were processed within two hours of endoscopic sampling.

### Single cell and nuclei preparation from fresh biopsies

Single cell suspension from mucosal biopsies were prepared fresh (modified from previous study (*6*). At least one biopsy from each sampling site was processed for single cells (sc), and the remaining tissue was fixed in 10% neutral buffered formalin at room temperature for 24 hours before embedding in paraffin (FFPE). Tissue dissociation is detailed in the online protocol (https://dx.doi.org/10.17504/protocols.io.bmdbk22n). Sample meta data and quality control were curated following BRISQ guideline.

### High throughput scRNA-Seq

Single cells were processed through NextGEM Single Cell Platform per manufacturer’s guide (10x Genomics, CG000204_ChromiumNextGEMSingleCell_3’ v3.1). The sequencing depth exceeded 30,000 reads per cell.

### FFPE multiplexed protein immunofluorescence (IF) and analysis

Tissue sections were processed and stained (*59*), with validated antibodies in Table S6. IF in Tissue sections were imaged by Leica SP8 or Leica STELLARIS confocal microscope with 40x lens at a pixel size 250 x 250 nm. The images were processed by *Cellpose* (*60*) for cell segmentation and counting. The threshold for positive staining was set at 30 with model “cyto”. The nuclear staining was counted using model “nuclei”.

### Spatial transcriptomics and analysis

Four TI FFPE blocks were processed per manufacturer’s guide for Xenium (10x Genomics, Xenium In Situ Gene Expression v1.0, CG000584 Xenium Analyzer). Human colon panel, customized probe panel (Table S6) and Cell Segmentation Staining Reagents Kit (10x Genomics, CG000749) were used to profile two 5 μm serial sections from each block. Cell segmentation was automated by Xenium Analyzer (10x Genomics). The cellular-level quantification was integrated and analyzed by *Seurat,* then the domain/niche analysis was conducted using *Banksy* and *FNN* (*61–63*) (supplemental method).

### Sequencing data preprocessing, integration and analysis

ScRNA-seq data was processed following a pipeline by *Seurat* with customized thresholding. Cell types were annotated based on marker genes and overall transcriptomes (*4–9*).

Differential expression genes (DEGs) on cell subpopulations and sample groups were extracted by Wilcoxon and linear-mixed models with set significance (adj. p-val < .05). Gene set enrichment analysis (GSEA) was performed on DEGs against major functional categories (supplemental method).

*Imputing cell differentiation* Macrophage trajectory was analyzed by *Monocle 2* (*64*). To characterize EC and FAE, transcription factor (TF) activities were estimated using *decoupleR* (*65*) (supplemental method).

*Imputing cell-cell interaction from ligand-receptor analysis* Ligand-receptor (LR) interactions were inferred for all cell types, pairwise in each sample using *CellPhoneDB* (*66*) (supplemental method).

### Disease scoring and network analysis

*Sample scores based on the expression of differentially expressed and IBD risk genes* Three sets of genes were used to quantify inflammation: 1) *InfDEG* - differential expressed genes on epithelial and immune lineages, 2) risk genes in genome wide associated studies (GWAS) loci (*29*) and 3) *UCSig* - inflammatory genes in UC (*6*). The inflammation score was assigned with the mean expression of genes in each sample (supplemental method).

*Co-expression analysis and samples scores based on key genes reflecting disease activity* We aimed to capture the pairwise gene-gene associations within lineage in each sample, where granularity of inflammation activity could be preserved. We proposed a framework utilizing this co-expression to construct networks and predict inflammation gene signatures (*CoInfSig*). The robustness of *CoInfSig* was assessed for generalization (supplemental method).

## ACKNOWLEDGEMENT

We are grateful to all the donors who consented to contribute samples to this study. We are grateful to Drs. Matthew Stephens, Sebastian Pott, Preety Bajwa for helpful discussions and to Drs. David Rubin, Russell Cohen, Sushila Dalal, Atsushi Sakuraba, Neil Sengupta, Dejan Micic, Edwin McDonald for clinical classification and tissue collection. We thank Sarbani Adhikari, Max Loy for clinical data collection, and the clinical coordinator team in Dr. Eugene Chang’s lab for their effort in obtaining patient consent. All tissue samples were from the University of Chicago Digestive Diseases Research Core Center, Center for Interdisciplinary Study of Inflammatory Intestinal Disorders (NIDDK P30 DK042086). Next-Generation sequencing was performed at the University of Chicago Functional Genomics Facility and computational resources were provided by the University of Chicago Research Computing Center. We also acknowledge the use of University of Chicago Human Tissue Resource Center, Integrated Light Microscopy Core and Cytometry and Antibody Technology Core (Cancer Center Support Grant P30CA014599), Integrative Clinical and Biospecimen Core and Northwestern University Metabolomics Core Facility for this study. This work is part of human Gut Cell Atlas Crohn’s Disease Consortium (https://helmsleytrust.org/gut-cell-atlas/).

## Data and materials availability

The FASTQ files can be accessed at Gene Expression Omnibus repository: accession code GSE266616 (token: qzgnwsyezpqjxeh). The integrated data matrices of single cell and spatial transcriptomes can be accessed at Zenodo: 10.5281/zenodo.22131807. The code for co-expression network analysis is accessible at (https://github.com/bingqing-Xie/CoSig/tree/main). Further information and requests for resources should be directed to and will be fulfilled by the lead contact, Anindita Basu. Request of preserved sample specimen will be fulfilled following the university regulations which will include, but not limited to materials transfer agreements.

## FIGURES AND FIGURE LEGENDS

Figure acronyms: IBD – inflammatory bowel disease; CD-Crohn’s Disease; IF – immuno fluorescence; FISH – fluorescence in situ hybridization; UMAP – uniform manifold approximation and projection; DDR tree – discriminative dimensionality reduction via learning a tree; DEG – differential expressed genes ; LR –ligand receptor; GSEA– gene set enrichment analysis;.

## SUPPLEMENTAL TEXTS

List of materials

Fig S1 to S5 Supplemental Methods

Data file Table S1 to S6 (Excel files)

Table S1 Sample meta data.xlsx

Table S2 Immune DEG and GSEA.xlsx

Table S3 Cell-cell interaction by Ligand-Receptor.xlsx

Table S4 Epithelial DEG and GSEA.xlsx

Table S5 Co-expression network.xlsx

Table S6 Antibody RNA-probe panels.xlsx

Supplemental References (*67–88*)

Supplemental figure acronyms: TI – terminal ileum; AC – ascending colon. Ctrl – control; Uni – noninflamed; Adj – adjacent to inflamed; Inf – inflamed; IF – immunofluorescence staining; GSEA – gene set enrichment analysis; LR – ligand-receptor. LND – *LCN2, NOS2, DUOX2*. H&E – hematoxylin and eosin; FAE – follicle associated enterocytes; G-like cell – Gastric chief- like cell; L – ligand; R – receptor. *CoInfSig* – inflammation signature genes from co-expression networks; HOSVD – higher-order singular value decomposition; TC – tensor component. *UCSig* – inflammatory signature genes from ulcerative colitis.

## Supplemental Figures

**Supplemental Figure S1.**
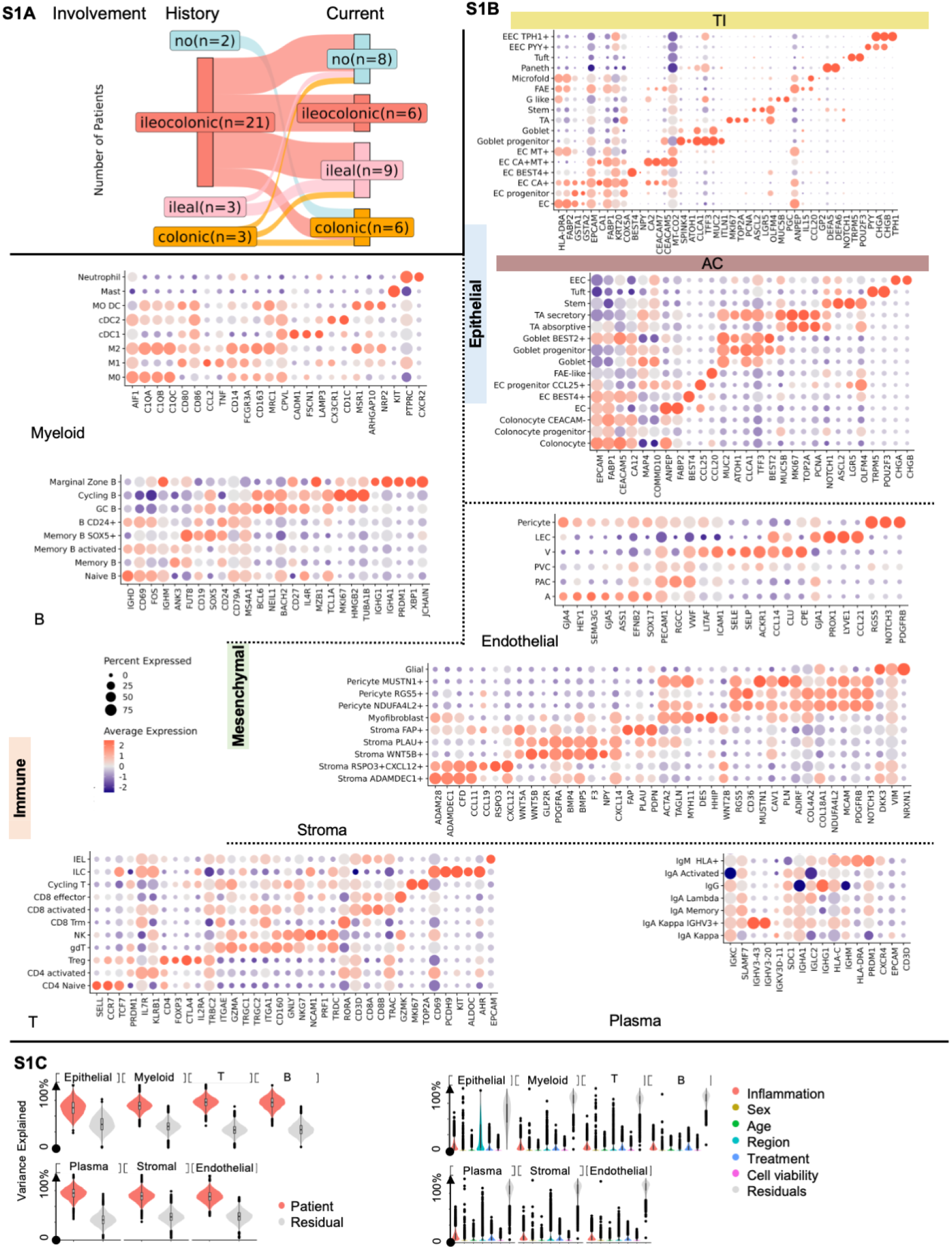
Biopsy dissociation assessment, cell type annotation and transcriptome variance analysis. A. Regional involvement shifts over time in Crohn’s Disease donors. B. Cell type annotation using gene markers. C. Assessment of transcriptome variance contributions. Gene expression aggregated by compartment per sample. A weighted linear mixed model was applied to quantify the contributions.

**Supplemental Figure S2.**
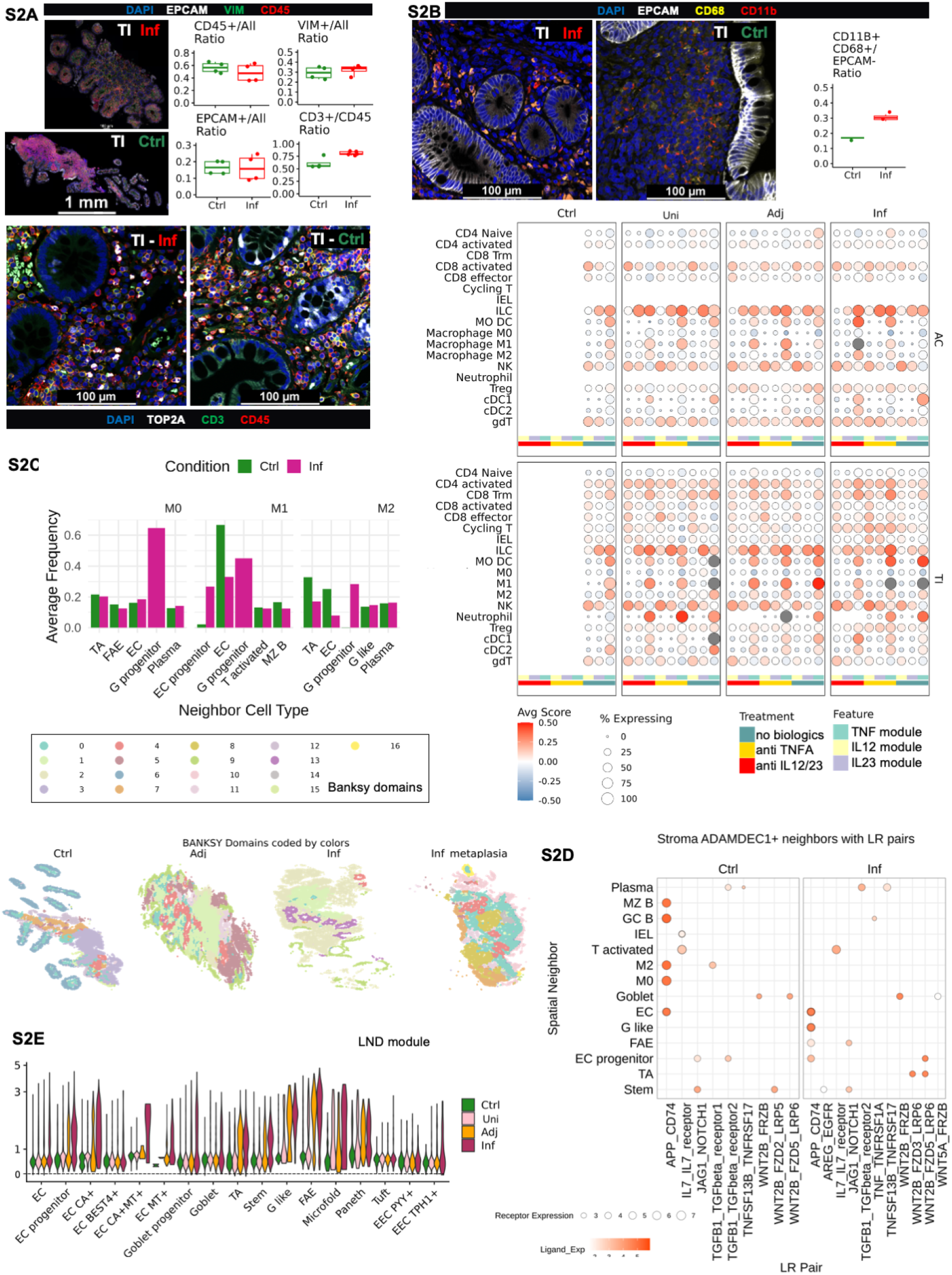
Changes in lineage composition, macrophage function and cellular neighborhood with inflammation. A. Lineage composition on **TI** tissue sections. Top left, IF images of **Ctrl** and **Inf TI** with lineage marker staining (EPCAM, CD45. VIM). Top right, lineage fractions per section in **Ctrl** and **Inf TI** sections; T cell fraction shown separately. Bottom, IF images of **Ctrl** and **Inf TI** with T cell marker staining (CD3, CD45) and proliferation marker (TOP2A). B. Macrophages in **TI** tissues. Top left, IF images of **Ctrl** and **Inf TI** with macrophage marker (CD68, CD11b) and epithelial marker (EPCAM). Top right, macrophage fractions in **Ctrl** and **Inf TI** sections. Bottom, cytokine modules (*TNF*, *IL12*, *IL23)* in immune cells across inflammation classification and biologics. Expression modules represent gene expressions associated with each cytokine. C. Cell neighborhood analysis based on spatial transcriptomics. Top, leading macrophage neighbors across domains, in Ctrl and Inf TI; Bottom, domain analysis across TI tissues. D. Stromal cell interaction in TI tissues. Spatial transcriptomes for stroma ADADEC1+ cell: X-axis, neighboring cells; y-axis, LR pairs. E. LND module expression in **TI** epithelial cells across inflammation classification. Violin plot; y axis, log transformed module scores.

**Supplemental Figure S3.**
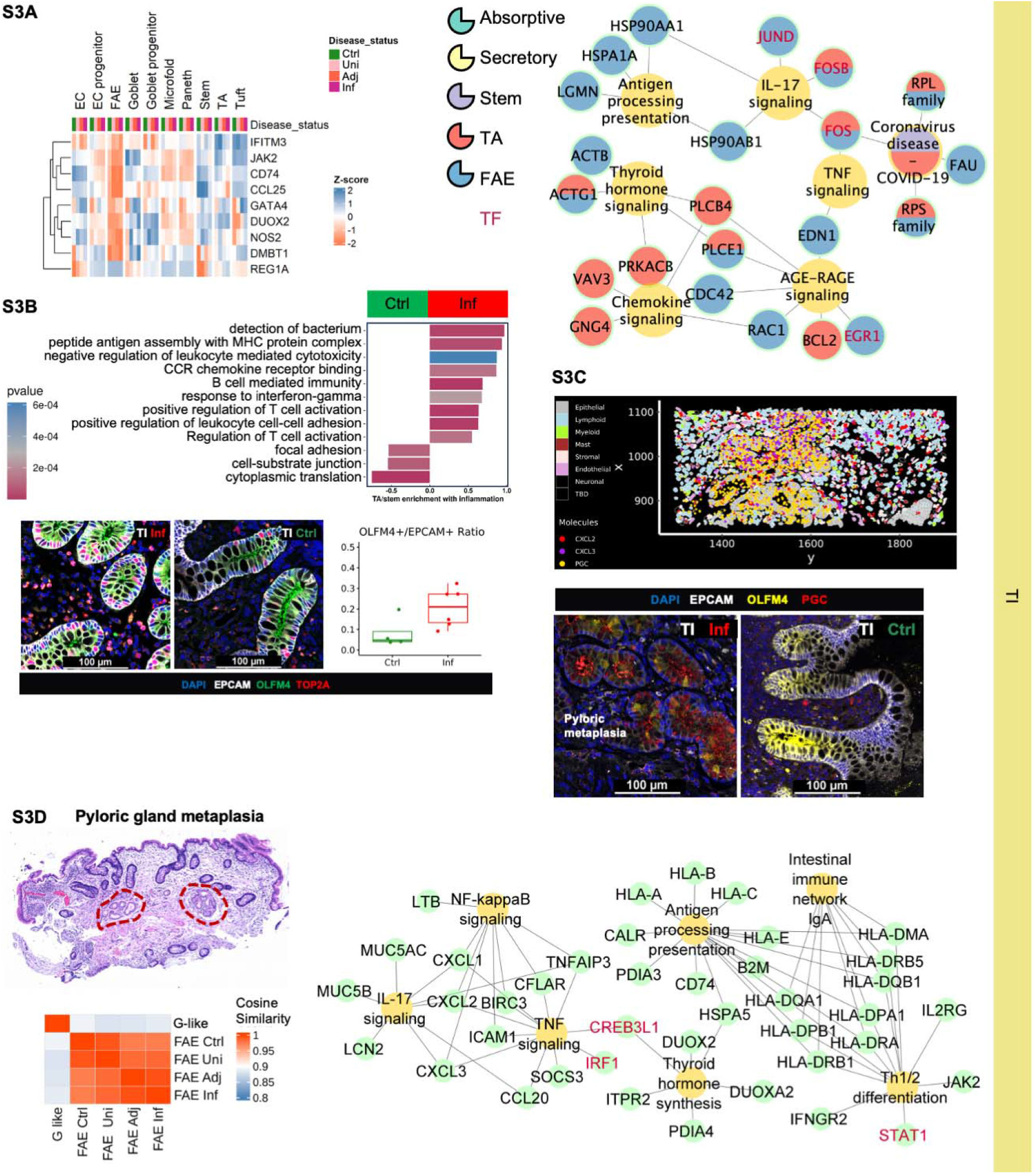
Epithelial cell composition and function in TI. A. Inflammatory gene programs in TI epithelial cells. Left, DEG across inflammation classification. Right, GSEA in **Ctrl TI**. Central node, enriched function; peripheral nodes, associated DEGs; Node and wedge colors indicate cell type; wedge area indicate cell type contribution. B. Stem and TA composition and functions from **Inf TI**. Top, GSEA in **Ctrl** and **Inf TI**. Bottom, IF – TA and stem in epithelial crypts and composition change from **Ctrl** and **Inf TI**. Marker – EPCAM (epithelial), OLFM4+TOP2A-(stem), OLFM4+TOP2A+ (TA). C. G-like cell in pyloric gland metaplasia in **Inf TI**. Left, Xenium images; cell colored by compartments; puncta by genes: PGC (G-like), CXCL2, CXCL3 (chemokines). Right, IF staining with EPCAM (epithelial), OLFM4 (stem and TA) and PGC (G-like). D. G-like cells location and function in Inf TI. Left top, H&E staining with metaplasia; left bottom, transcriptomic comparison between G-like cell and FAE. Right, GSEA of G-like cells. Central node, enriched, peripheral node, associated gene.

**Supplemental Figure S4.**
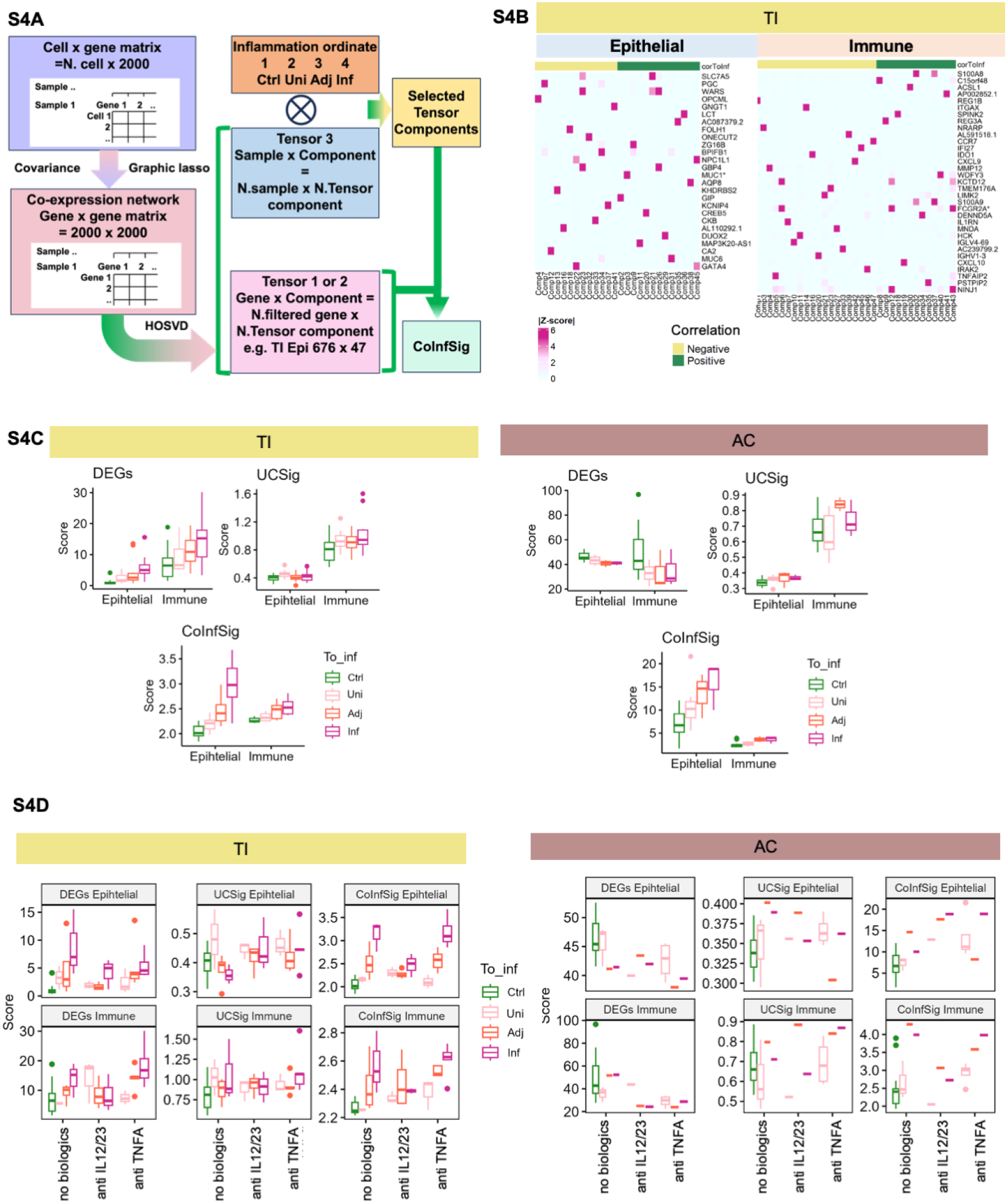
Co-expression analysis workflow and findings in **TI**. A. Framework for extracting *CoInfSig* from co-expression networks using HOSVD. B. Example HOSVD loading matrices for top loading genes in TCs in **TI** epithelium. C. Sample scores derived from DEGs, UCSig, *CoInfSig.* Left, TI; Right, AC. D. Sample scores by treatment group across inflammation classifications. Left, TI. Right, AC.

**Supplemental Figure S5.**
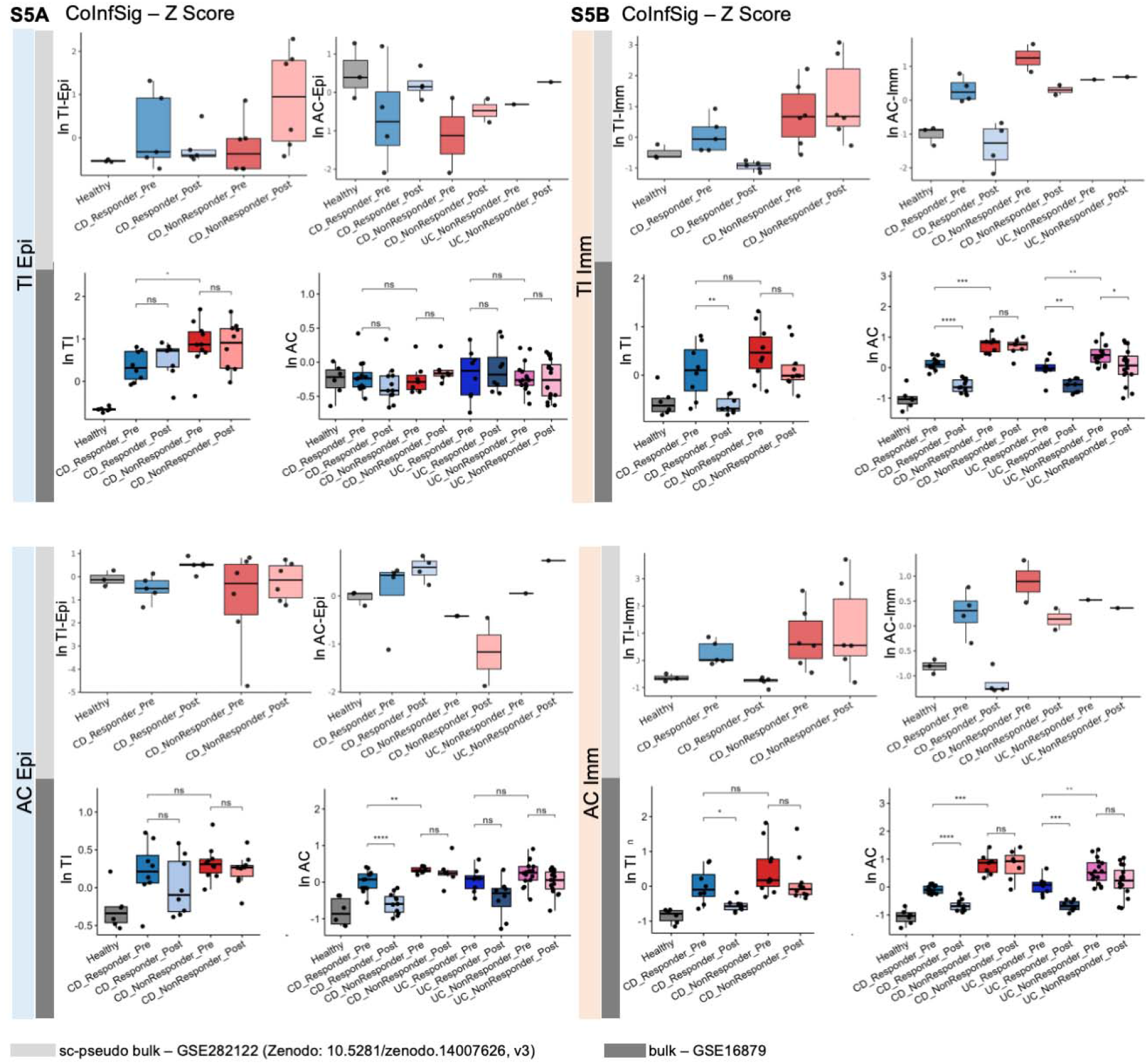
Applications of co-expression markers in external data sets. A. Robust epithelial *CoInfSig* Z-score in external data with longitudinal sampling in CD anti-TNFα responders and non-responders. Top, TI epithelial *CoInfSig*; Bottom, AC epithelial *CoInfSig*. B. Robust immune *CoInfSig* Z-score in external data with longitudinal sampling in CD anti-TNFα responder and non-responder. Top, TI immune *CoInfSig*; Right, AC immune *CoInfSig*. ns, not significant; *, *p* <0.05; **, *p* <0.01; ***, *p* <0.005; ****, *p* <0.001. Reused data: scRNA-seq – GSE282122 (10.528/zenodo.14007626); bulk RNA-seq – GSE16879.

## Supplemental Methods

### Data alignment and pre-processing

*CellRanger* (10x Genomics, 7.0.1) was used to align the fastq reads to the reference genome (*67*) (gencode hg38 v.32). Digital gene expression (DGE) matrices were generated from sequence reads mapped to both exonic and intronic regions in the human genome. To select high-quality cells, we set quality control cutoffs 1) unique molecular identifiers (nUMI) > 1000; 2) number of genes between 200 – 6000; 3) the percentage of mitochondrial reads (MT%) < 50% 4) Scrublet (*68*) doublet scoring (ds) < 0.2. The filtered DGE matrixes (36,601 genes across filtered cells) were used for downstream analysis. We followed a standard preprocessing pipeline using the single-cell analysis suite, *Seurat* v4.1.1 (*69*). A logarithmic normalization (log() + 1) method was applied to transfer gene expression counts, scaled by 10,000 (TP10K). The gene expression matrix was centered, and nUMI and MT% were regressed out using the *ScaleData* function.

### Batch correction, cell clustering, annotation and composition analysis

The top 2,000 highly variable genes from the integrated data matrix were selected for principal component analysis to generate 100 principal components (PC). The ‘Patient’ variable was used for *Harmony* (*70*) on the PC to remove donor-specific variance. We selected the top 50 PC, based on their variance explained. The selected PC were used in further exploration of the data, such as UMAP dimension reduction, construction of K-nearest neighbor graph, and shared nearest neighbor modularity optimization-based clustering (*71*). We used UMAP to project the cell population for 2D visualization. Hierarchical clustering on the shared nearest neighbor graph was used to infer the cluster structure of the cells; the ‘resolution’ was set between 0.1 – 0.8 to build multi-resolution clusters. Cluster marker genes were extracted by *FindMarkers* using the Wilcoxon Rank Sum test (*72*). The clusters were first annotated at lineage and compartment levels, such as epithelium, immune (including T, B, Plasma, and Myeloid cells), and mesenchyme (including stroma and endothelium) by cross-referencing the resulting clusters with known cell-type markers. At this level, TI and AC were separated for region specificity in the epithelium.

The compartments were then isolated and followed by subsequent processing to identify fine-grain cell type clusters. Labels transferred from published human gut single-cell datasets (*4–7, 73*) and marker genes were used to annotate cell types, with prior knowledge of human intestines. Cell types in each compartment were displayed in UMAP, and the cell types belonging to a different compartment biologically were removed from the UMAP for visualization. Cell counts were normalized by the population count (epithelium, immune, and mesenchyme). The difference of mean for cell ratios regarding multiple variables, such as inflammation (**Inf** vs. **Ctrl**), were tested using the Wilcoxon Rank Sum test and p values were corrected by false discovery rate (FDR). Correlation between cell abundance and inflammation classification was assessed using *cor_test* in *ggpubr* (*74*).

### Differential expression analysis on multiple comparison groups

To investigate the difference in region, disease, and inflammation, we performed differential expression (DEG) analysis by *FindMarkers* in *Seurat* using the Wilcoxon Rank Sum test, MAST (*75*) with fixed effects, and Dreamlet (*76*) for the pseudo-bulk with mixed effects on each cell type. The statistically significant (adjusted p-value, adj. p-val. < 0.05) markers were extracted for each cell type and its contrast groups. Shared genes from both Wilcoxon rank-sum test and pseudo-bulk tests on the ‘control vs. inflamed’ contrast group were used as robust inflammation signatures. Gene set enrichment analysis (GSEA) on the log fold changes of genes was performed using R library *ClusterProfiler* (*77*) against the Gene Ontology (GO) terms and the KEGG (*78*) pathways. Visualization of the dot plot, violin plot, heatmap, upset plot, and enrichment plot was acquired by *ggplot2* (*79*), *Seurat*, *enrichPlot* (*80*), and *ComplexHeatmap* (*81*). For TI epithelial, we set several groups, Stem (stem, TA), absorptive (EC-enterocyte, EC BEST4+, EC CA+, EC progenitor), and secretory (goblet progenitor, goblet, EEC TPH+, and EEC PYY+), and FAE (FAE follicle associated enterocyte). Intersection of DEG for all cell types within each group was noted as conserved markers. The illustration between enriched functions/pathways and genes was plotted using Cytoscape 3.8.2 (*82*) with yFiles (*83*) organic layout. Color of each node was coded for the cell type where the DEG was identified.

### Imputing cell differentiation

Macrophage trajectory was built using macrophage M0, M1 and M2 from myeloid compartment. FAE trajectory was built with FAE, EC, EC progenitor in TI (up to 2000 cells per inflammation classification for each cell type). After sampling the selected cells, data was preprocessed using *Seurat* v4.1.1 pipeline. The cell ordering for pseudo-time and trajectory estimation was performed by *Monocle 2* (*64*) using top 2000 highly variable genes that segregate cells by transcriptomic similarity. The trajectory was reduced to 2D space using *DDRtree*. Cells were ordered with a given starting cell type (e.g. M0 for macrophage). Branch driving genes (genes that are differentially expressed between branches) were extracted by *BEAM*. Trajectories and heatmaps were plotted using the *plot* function in *Monocle 2*. Boxplot for cellular state ratio was plotted using *ggplot2*.

To further characterize EC and FAE, transcription factor (TF) activities were estimated using *decoupleR* (*65*) for these two cell types. Active TF were calculated using *run_ulm,* of which the top 100 most variable TF were selected across all inflammation classification of EC and FAE. TF were further filtered for positive expression in EC and FAE single-cell transcriptomes. Heatmap for Z-scores was used for differential TF activity visualization between EC and FAE.

### Imputing cell-cell interaction from ligand-receptor analysis

Ligand-receptor (LR) interactions were inferred for all cell subtypes, pairwise in each sample using CellPhoneDB (*66*, version 4). The predicted LR interactions were binarized where LR interactions without statistical significance (p ≥ 0.05) were removed. The LR interactions were aggregated and averaged within each sample, then grouped by variables of interest, such as Inf, and Ctrl. Within each group for any given LR pair, we used the sample count with this LR interaction over the total number of samples as the normalized LR count. For a given inflammation group (e.g. **Inf**), we scored any pair of cell types by accumulating the normalized LR counts over all LR pair counts, to construct a weighted cell-type wise LR network. The top cell pairs by LR were visualized by *Cytoscape* using *yFiles* organic layout. We colored the cell type by their lineage annotation and the edge thickness was used to indicate the weight. The total normalized count of LR interactions among compartments was calculated to dissect highly interactive compartment pairs. The cellular input in the interaction network is calculated for cell-specific LR weighted counts/Total LR weighted counts in the network.

### Spatial transcriptome analysis

Cell segmentation was automated based on the cellular staining from the segmentation add-on by Xenium Analyzer (10x Genomics). Output data was imported and integrated using *Seurat*, and segmented cells were selected using the cut off: count of probe puncta (*nCount_Xenium*) >10. We normalized and scaled the integrated data using *SCTransform* in *Seurat.* Top 418 variable genes were selected for PC analysis. No tissue specific variance was removed. We selected the top 30 PC to further explore the data, including K-nearest neighbor graph construction, shared nearest neighbor modularity optimization-based clustering, and UMAP dimension reduction. The resolution for hierarchical clustering on the shared nearest neighbor graph was set between 0.1 – 0.4. Cluster marker genes were extracted by *FindMarkers* using the Wilcoxon Rank Sum test. The clusters were initially annotated at lineage level (epithelium, immune and mesenchyme) by lineage-specific genes. Each lineage was subsequently split and processed to identify cell type clusters. Marker genes from these cell type clusters were used for annotation, leveraging cell type marker genes from sc transcriptome. Annotation strategy for sc transcriptome was applied to the spatial transcriptome, except for one cluster—secretory TA that shared less gene markers with the counterpart cell type in sc transcriptome. We used UMAP to visualize the cells in epithelial, immune and mesenchymal lineages across all samples. Across each tissue sections, we utilized *Banksy* to identify cellular domain/niches at various sizes, we used *FNN* to identify cellular neighbors with the proximity (*62,63*). We selected cell pairs with the top normalized frequency from each cellular domain and constructed the spatial cell interaction network. LR genes were referenced to the databased in *CellPhoneDB* (*66*).

### Disease scoring and network analysis

To quantify inflammation, we relied on the mucosal inflammation classification by endoscopy and extracted cell-level transcript expressions, including those in IBD Genome-Wide Association Study (GWAS) (*29*) loci.

### Sample scoring based on the expression of lists of genes

From the inflammation classification, we first established the positive inflammation signature for the two extreme classifications, **Ctrl** and **Inf**, by DEG analysis on epithelial and immune lineages split by region. Most shared significant DEG (FDR-adjusted 0.05, common to majority if not all of the cell types) upregulated in **Inf** among the cell types within each lineage were retrieved and merged on lineage level as the DEG signature. For epithelial lineage in terminal ileum (TI), the *InfDEG* are *S100A11, MUC1, PIGR, CHRM3, REG1B, REG1A, CCL20, ALCAM, MECOM, KCNIP4, CCL28, ATP10B, CFB, HLA-DPA1, HLA-DPB1, AGR2, NPC1L1, GATA4, LYN, AC083837.1, LCN2, DMBT1, IFITM3, FOLH1, CCND2, LYZ, OLFM4, GPX2, IFI27, IGHA1, DUOX2, NOS2, ONECUT2, TSHZ2, SLC40A1, and ITLN1.* For epithelial lineage in ascending colon (AC), the *InfDEG* are *BRINP3* and *FAM3B.* For immune lineage in TI, the *InfDEG* are *GBP1, STAT1, WARS, S100A9, GBP5, ACSL1, IRF1, GK, S100A8, SOD2, TAP1, TYMP, GBP2, ISG20, ANKRD22, LIMK2, LAP3, AQP9, SNX10, REG1A, GCH1, SOCS3, APOL6, PDE4B, IFITM3, MX2, SERPINA1, FCGR2A, HIF1A, CD274, VCAN, GLIPR2, FNDC3B, TIMP1, CD300E, FCGR3A, MXD1, DENND5A, IFITM2, PARP9, GBP4, MX1, MYOF, TNFRSF1B, FCN1, LILRA5, CDKN1A, KYNU, C15orf48, NLRC5, TNFSF13B, RBMS1, LILRB3, G0S2, HIVEP2, MYO1G, ITGAX, IFIT3, EHD1, FGR, DDX60L, AC083837.1, SLC16A10, BASP1, ATP13A3, JAK3, RIPK2, AL157871.4, LMNB1, STK10, PPIF, TNFSF10, MNDA, SLC25A37, NCF2, IGLC1, EREG, INHBA, B4GALT5, IQSEC1, SAMD9L, STAT4, CALHM6, CUL1, CSF2RB, ELL2, SLC7A5, AC016831.7, MEFV, IL7R, CD44, AREG, RIPOR2, PIGR, MOB3B, TREM1, FLT1, PSTPIP2, LRRK2, PLAC8, MAP3K20, LINC01619, MAP4K4, DDX58, SELL, TFRC, PRKCB, ECE1, C5AR1, PLCB1, CPD, AC099489.1, IFI27, XBP1, SAT1, IGLV3-1, DCC, NFKBIA, SQSTM1, IGLC3, IGKC, IGHA1, IGHM, IGLC2, GZMB, and IFNG. For immune in AC, the InfDEG are CD48, EEF1A1, IGHA2, JCHAIN, MT-CO1, RPL8, RPL9, RPS14, RPS26, RPS3, SPINK2, TIMP1,* and *XIST.* Significant DEG on all cell types within each lineage were used as the first set of inflammation signature genes. We also included previously reported inflammation-related genes in IBD-UC (*6,84*) as contrast sets. Using the genes obtained from pseudo-bulk analysis, each sample was either averaged into a single vector representing the gene expression for all cells or aggregated into multiple lineage-specific vectors (e.g., sample 1: epithelial, sample 1: immune, etc.) where the gene expression was averaged within the lineage. The inflammation score was then computed from the mean expression value of the corresponding inflammation signatures for each lineage.

In addition to analyzing the inflammation activities by individual genes (e.g. DE analysis), we aimed to capture the pairwise gene-gene associations within epithelial and immune lineage within each sample, where heterogeneity among individuals could be preserved. The gene-gene associations were inferred from gene co-expression patterns, where the correlation of expression levels for any pair of genes was measured among a group of cells. Genes used to build the co-expression matrix were top 2000 variable genes from the vst selection in Seurat (in the epithelial and immune lineages across all samples respectively), with log normalized expression over 0.1.A positively correlated gene-pair indicates that the gene expressions increase or decrease simultaneously among a group of cells. A negatively correlated gene pair suggests that the gene expressions vary in the opposite directions. Alterations of gene-gene association, e.g., genes were co-expressed only under certain conditions, and shift of hub genes (central nodes in the association network that are responsible for the connectivity of the network) were computed as follows: We first computed the pairwise gene co-variance matrix for each sample within one lineage: epithelial, or immune, where the co-variance for any pair of gene expression vectors *x, y x, y* is,

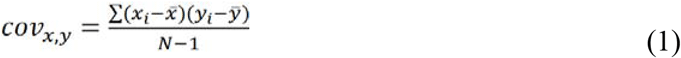

*NN* is the number of cells and *x̄x̄* is the mean of *xx*. A co-expression network was constructed with graphical lasso using glasso (*85*) on the gene covariance matrix from each sample to prune false positive connections. We used matrix representation to preserve the structure of the network, avoiding its reduction into pairwise tuples. To capture inflammation variation among samples, we included an additional matrix dimension to encode sample ID and inflammation classification. The co-expression networks for all samples formed a 3D matrix (or mathematically, a tensor, we will use this notion throughout the rest of the section) *X∈ℝ^G×G×S^ X∈ℝ^G×G×S^*, where *GG* is the set of highly variable and expressed genes (previously described), *SS* is the sample set, and *RR* is the tensor component set. High order singular value decomposition (HOSVD) using CANDECOMP/PARAFRAC (CP) method implemented in R package *rTensor* (*86,87*) was adopted to identify structure variations along all axes in the 3D tensor *XX* with *RR* components. The output contains a 3D diagonal tensor ∧*∈ℝ^R×R×R^* ∧*∈ℝ ^R×R×R^*, and three projection matrices, *U*^1^∈ℝ*^G×R^*U^1^∈ℝ*^G×R^*, *U*^2^∈ℝ*^G×R^*U^2^∈ℝ*^G×R^* and *U*^3^∈ℝ*^G×R^*U^3^∈ℝ*^G×R^*. The three projection matrices contain two highly homogeneous gene projections and one sample projection. The 3D tensor *XX* is approximated by,

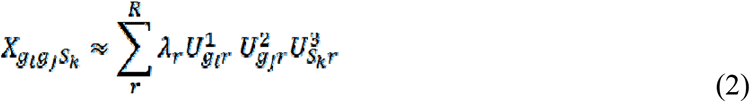

where *g_i,_ g_j_∈G,S_K_g_i,,_g_j_∈G, S_K_∈S∈S, r∈Rr∈R, λ_r_λ_r_,* is the *rr^th^* entry in cubical diagonal tensor ∧∧, 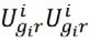 is the rank-one tensor for gene *g_i_g_i_* and *rr* in orthogonal matrix *U^i^U^i^*. We selected the tensor components based on Spearman correlation (*p < 0.05p < 0.05)* between the sample projections and inflammation index. The genes with significant projection values (*Z>2*) in the selected tensor components were identified as the driver genes of those inflammation-related components and denoted as co-expressed using its direct (first order) neighbor from the co-expression network. Note that each edge in the co-expression network indicates a co-occurrence pattern for a gene pair. When aggregating multiple co-expression networks, we defined an edge prevalence ratio score *er er*to measure the occurrence of a gene-gene association among a set of samples,

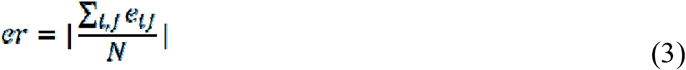

where *e_ij_* = 1*e_ij_* = 1if a positive edge *ii* exists in sample *jj*, *e_ij_ = -1e_ij_ = - 1*if a negative edge *ii* exists in sample *jj*, and *NN* is the number of samples within the sample set. A threshold on edge prevalence ratio (|*er*| > 0.3|*er| > 0.3*) was applied to detect homogeneous/common response (i.e., shared by multiple samples within a group) to the inflammation in CD, while maintaining some sample-specific responses. The resulting networks from the inflammation classification illustrate the inflammation-related alteration on gene-gene co-expression. The centrality properties betweenness and closeness (88) were used to evaluate the network, and the definition is as follows: Given *g_i,_ g_j_*, *g_k_ ∈ G g_i,_ g_j_*, *g_k_ ∈ G*, the betweenness is

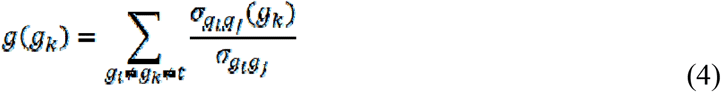

where *σ_gigj_σ_gigj_* is the total number of shortest paths from gene *g_i_g_i_g_i_* to gene *g_j_g_j_g_j_* and *σ_gigj_*(*gk*) *σ_gigj_*(*gk*) *σ_gigj_*(*gk*) is the number of paths that pass gene *g_k_g_k_g_k_*. Given *g_i_g_j_* ∈ *Gg_i_g_j_* ∈ *Gg_i_g_j_* ∈*G*, the closeness^33-35^ is

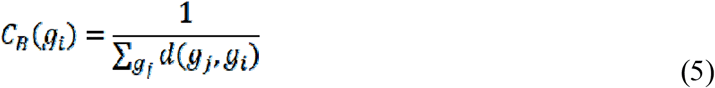

where *d(g_j,_ g_i_)d(g_j,_ g_i_)* is the length of the shortest path between gene *g_i_g_i_* to gene *g_j_g_j_*. The network centrality metrics were computed by Cytoscape.

## Randomization analysis on sample size and network-based inflammation signatures

To assess the signature rigor and patient heterogeneity on CoInfSig, we conducted randomized sampling by drawing a fixed sample size α from each inflammation group for both epithelial and immune lineages. We varied α from 3 to 14 (the maximum sample size within any inflammation group; we allowed duplicated samples as replacement if sample size was smaller than α) and for each α, we permuted the randomized sampling 100 times. We performed the HOSVD analysis and inflammation signature selection process for all random sets. We then computed the probability P_g_ for any gene g: P_g_ =I_g_ /100, where I_g_ is the number of times that the gene g was selected as a CoInfSig element. The probability for the CoInfSig with all samples present was visualized as a heatmap. The columns were ordered by the smallest sample size for which a signature has > 50% probability to be a stable signature, given the sample size.

## Sample scoring based on the expression of selected marker genes

To investigate the usability of these marker genes in differentiating biologic response in CD patients, we applied Z-scoring based on the expression levels across external data sets: for sc transcriptomes (*50*), the pseudo bulk transcriptomes were aggregated from all the cells within one lineage (epithelial or immune), and expression level for each marker gene were calculated after normalization of counts by sample-lineage cell counts and sequencing depth; for bulk transcriptomes (*51*), TPM were used to calculate the normalized gene expression. Sample Z-scores were calculated by averaging the Z-scores in each CoInfSig marker sets. Welch t-Test was used to compare scores between two groups.

